# The molecular consequences of androgen activity in the human breast

**DOI:** 10.1101/2022.04.22.489095

**Authors:** F. Raths, M. Karimzadeh, N. Ing, A. Martinez, Y. Qu, T.Y. Lee, B. Mulligan, S. Devkota, B. Wang, A.E. Giuliano, S. Bose, H. Goodarzi, E.C. Ray, X. Cui, S.R.V. Knott

## Abstract

The mammary gland has been extensively studied for estrogen and progesterone reactivity, but the molecular effects of androgen in the breast remain largely unexplored. Transgender men are recorded female at birth but identify as male and may undergo gender-affirming androgen therapy to align their physical characteristics and gender identity. Here we perform single cell resolution transcriptome, chromatin, and spatial profiling of androgen treated breasts from transgender men. We find male-biased androgen receptor gene targets are upregulated in cells expressing androgen receptor, and that paracrine signaling drives sex-relevant changes in other cell types. We observe an altered epithelium, shifts in immune populations, and a reduction of capillary vasculature. Finally, we find evidence of the metabolic impact of androgen and identify a gene regulatory network driving androgen-directed fat loss. This work elucidates the molecular consequences of androgen in the human breast at single cell resolution.

## Introduction

The mammary gland relies on continuous hormonal modulation during development, menstrual cycles, pregnancy, and lactation. The most critical hormonal regulators are estrogen, progesterone, prolactin, and oxytocin^1–3^. Interestingly, only ∼10% of breast epithelial cells express hormone receptors, suggesting elaborate paracrine signaling cascades to communicate growth and secretory stimuli to the tissue^4^. The androgen receptor (AR) is also represented in the breast, and it is generally accepted that androgens counteract the effects of estrogen and progesterone and can therefore inhibit thelarche (pubertal breast development)^5, 6^. The precise molecular functions of androgen in the normal breast, however, have been widely ignored, while its implications for breast cancer progression have long been debated. AR is expressed in 60-90% of breast cancers, and recent data shows that AR is in fact a tumor suppressor in estrogen receptor positive (ER^+^) breast cancer, which opens new pathways for androgen targeted cancer treatments and demands a deeper understanding of the molecular dynamics of androgen action on the various cell types of the normal breast^7, 8^.

While murine models have been instrumental for our understanding of breast cancer biology, they pose considerable limitations for the study of healthy tissue. This is due to differences in human and murine hormonal cycles, mammary gland development and distribution of hormone responsive cell types^9^. One opportunity to study androgen signaling in humans lies in the analyses of tissues obtained from transgender men receiving gender-affirming care. Transgender is a term to describe a discordance between a person’s sense of gender identity and their sex recorded at birth^10^. Transgender men are individuals who were recorded female at birth, but identify as male, while cisgender women are recorded female at birth and identify as such. To address their physical discordance, many transgender men elect to undergo gender-affirming androgen therapy, which helps to align their physical characteristics with their gender identity by inducing a wide range of masculinizing physiological changes^11, 12^. Among the desired effects are increased facial and body hair, a deeper voice, termination of menstrual cycles, gain in muscle mass, body fat redistribution, and involution of breast tissue. Therefore, the treatment is often complemented by gender-affirming mastectomy^13, 14^. While the physiological effects of androgen therapy are well documented, the molecular and cellular implications of this treatment remain uncertain. This is due to a scarcity of high-quality transgender data and a general lack of understanding about the molecular functions of androgen in various hormone responsive tissues other than the prostate^15^.

The presented study aims to elucidate the androgen signaling landscape in healthy human breast tissue under long-term androgen exposure. We performed single nucleus RNA and ATAC (assay for transposase-accessible chromatin) sequencing on gender-affirming mastectomy specimens from transgender men undergoing androgen therapy and compared them to corresponding samples from cisgender women who had elective mammary surgery. We also used Co-Detection by Indexing (CODEX) multiplex immunohistochemistry (IHC) staining to describe changes in epithelial structure and cell-cell interactions. By integrating datasets of three pivotal modalities in the hierarchy of gene regulation and spatial distribution of cell types within mammary tissues, we aimed to deconvolute direct and indirect androgen targets, their modes of regulation, and the resulting phenotypic changes in the detected cell types. This multi-modal single cell atlas and the associated findings represent a comprehensive resource for studying the molecular consequences of androgen activity in the human breast.

## Results

### Single nuclei RNA and ATAC sequencing reveals breast cells are silenced by androgen exposure

To profile androgen-induced mammary gland alterations, we analyzed nuclei and tissue from breasts obtained from 9 transgender men who underwent gender-affirming androgen treatment and subcutaneous mastectomy. For controls, we studied corresponding breast tissue from 9 cisgender women who had elective mammary surgery (Fig. 1A, S1A-C). In this manuscript, for brevity, we refer to samples obtained from cisgender men, transgender men, and cisgender women as cis-male, trans-male, and cis-female respectively. In total, we analyzed 66,926 cis-female and 38,762 trans-male nuclei with snRNA-seq, as well as 27,459 cis-female and 30,927 trans-male nuclei with snATAC-seq. We classified cell types in the transcriptomic data from curated marker gene sets and then used their expression profiles to annotate corresponding snATAC-seq populations based on promoter accessibility (Fig. 1B, S1D, E, Table 1). In addition, we developed 8 tissue microarrays (TMA) from the left and right breasts of most of these individuals, which were analyzed using a custom CODEX antibody panel designed based on findings from the nuclei data (see Methods). This allowed us to further study 156,842 cis-female and 161,241 trans-male cells for their spatial context via image analysis (Fig. 1C, S1C).

**Figure 1:**
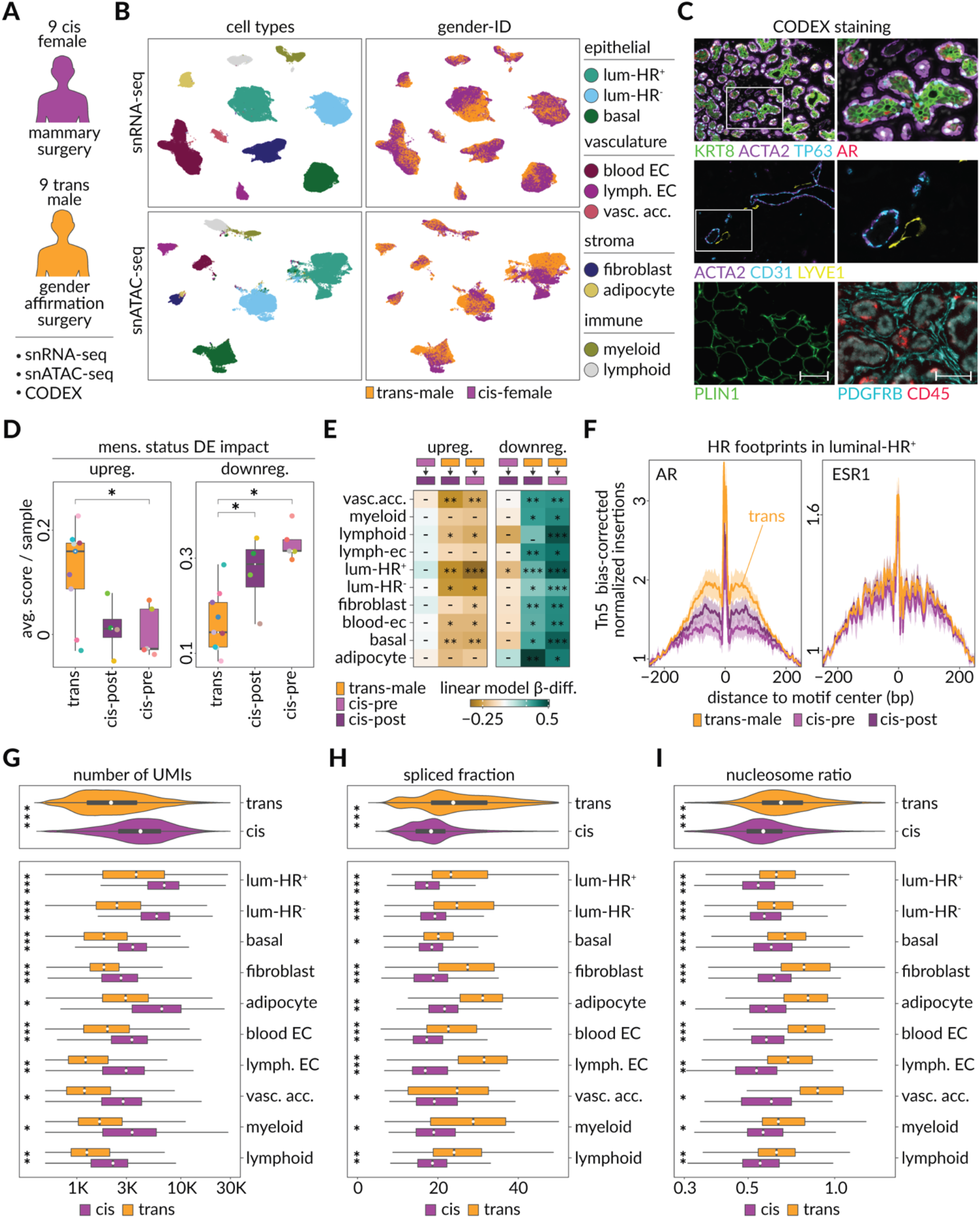
Multi-modal single nuclei sequencing, and spatial proteomics identify molecular distinctions between the breasts of transgender men and cisgender women. **A)** We performed single-nuclei RNA-seq (snRNA-seq), single-nuclei ATAC-seq (snATAC-seq), and multiplexed proteome tissue phenotyping (CODEX) on flash-frozen and FFPE embedded breast tissues from elective mammary surgeries in nine cisgender women (cis-female) and gender-affirmation surgeries in nine transgender men (trans-male). **B)** UMAP plots annotated by cell type (left) and gender-ID (right), with snRNA-seq on top (103,405 cells) and snATAC-seq data (49,728 cells) on bottom. (luminal-HR+ = hormone receptor expressing luminal cells, luminal-HR– = hormone receptor negative luminal cells, basal = basal/myoepithelial cells, blood EC = blood endothelial cells, lymph. EC = lymphatic endothelial cells, vasc. acc. = vascular accessory cells). **C)** Fluorescence microscopy images of the breast tissue with each marker identifying the major detected cell classes and structures: KRT8; pan-luminal, TP63; basal, AR; luminal-HR+, ACTA2; smooth muscle structures, CD31; endothelial, LYVE1; lymphatic vessels, PDGFRB; fibroblasts, CD45; immune, PLIN1; adipocyte. Scale bars: left column = 100 µm, right column = 50 µm. **D)** Boxplots showing average differential expression impact scores for each sample organized by menstrual status (trans-male = orange, pre-menopausal cis-female = light purple, post-menopausal cis-female = dark purple). Differentially expressed genes with an avg. log2FC > 0.5 (upregulated, p-values, Wilcoxon: trans-male vs. cis-pre = 0.029, trans-male vs. cis-post = 0.076, cis-pre vs. cis-post = 0.9) or < −0.5 (downregulated, p-values, Wilcoxon: trans-male vs. cis-pre = 0.001, trans-male vs. cis-post = 0.05, cis-pre vs. cis-post = 0.29) between trans-male and cis-female were used to score all cells within a sample, providing a metric on how much a sample contributes to the observed differential gene expression. **E)** Heatmaps show the effect size of a linear association between a cell’s differential expression impact score and the sample identity (trans-male, pre-menopausal cis-female, and post-menopausal cis-female). Each row used the cell within one cell type, and each column used two of the three sample identities. The perturbation signature of a cell uses upregulated genes (left panel) or downregulated genes (right panel) to score the cell’s transcriptome. Labels indicate p > 0.05 (–), p < 0.05 (*), p < 0.01 (**), or p < 0.001 (***). **F)** Motif footprints for Androgen Receptor (AR, AR-CisBP M03389_2.00) and Estrogen Receptor (ESR1, ESR1–CisBP M03356_2.00) in trans-male, pre-menopausal cis-female, and post-menopausal cis-female luminal-HR+ cells of the snATAC data. Horizontal axis indicates relative position to the center of the motif. Vertical axis indicates the transposase bias adjusted footprint signal. **G)** Number of unique molecule identifiers (UMIs) detected in each cell of the snRNA-seq data, split by cell types and gender-ID. Horizontal axis limits were set to 350-30,000, excluding 43 outliers to improve interpretability of the plot. (p-values, Wilcoxon: (***) p <= 5.29 x 10-203, (**) p <= 1.63 x 10-97, (*) p <= 2.66 x 10-71) **H)** Average fraction of spliced transcripts detected in trans-male and cis-female cells of the snRNA-seq data as calculated with velocyto (La Manno et. al., 2018). Horizontal axis limits were set to 0-50, excluding 49 outliers to improve interpretability of the plot. (p-values, Wilcoxon: (***) p <= 2.32 x 10-277, (**) p <= 6.33 x 10-93, (*) p <= 5.04 x 10-6) **I)** Degree of general chromatin condensation as the ratio of nucleosome bound to nucleosome free genomic fragments in each cell of the snATAC-seq data, split by cell types and gender-ID. Horizontal axis limits were set to 0.3-1.5, excluding 90 outliers to improve interpretability of the plots. (p-values, Wilcoxon: (***) p <= 3.76 x 10-137, (**) p <= 4.67 x 10-45, (*) p <= 7.64 x 10-5)

The epithelial compartment was composed of both hormone receptor positive and hormone receptor negative luminal groups (luminal-HR^+^ and luminal-HR^-^, respectively), as well as basal/myoepithelial cells. Within the vasculature, we observed blood- and lymphatic endothelial cells (blood EC and lymph. EC) as well as vascular accessory cells (vasc. acc.). Among the stromal populations we detected adipocytes and fibroblasts and in the immune compartment we found distinctive myeloid and lymphoid lineages. Except for the vascular accessory population, all groups were also present in the snATAC-seq libraries, with proportions showing significant correlation among samples (Fig. 1B, S1F, G). Using CODEX antibody staining patterns, we were also able to annotate most cells in the TMAs based on these categories. However, a subset of stromal cells was found to be negative for all markers (stroma-other) and a group of cells that were exclusively stained for ENAH could not be reliably identified (Fig. S1H, I).

Since our cohort consisted of pre- and post-menopausal cisgender women, we studied how age and menstrual status impacted these samples (Fig. S1J, Table 2). An examination of the distribution of globally up- and downregulated genes between transgender men and pre- and post-menopausal cisgender women, found that androgen treatment was the most significant driver of variation, while pre-menopausal samples appeared to show an overall stronger distinction compared to their post-menopausal counterparts (Fig. 1D). An analysis of individual cell types showed similar patterns and found differentially expressed genes had high correlation between pre- and post-menopausal samples (Fig. 1E, S2A, S3). Androgen therapy was also found to induce the most significant increase in chromatin accessibility when examining the binding for AR and ESR1 in luminal-HR^+^ cells (Fig. 1F). Interestingly, lowered estrogen levels in post-menopausal cisgender women had no discernable impact on ESR1 chromatin signal, but elevated accessibility for AR, presumably through loss of ESR1/AR antagonism (Fig. 1F)^16^.

A global unbiased analysis of our nuclei data further confirmed that androgen treatment was the primary factor of separation within the detected cell types, and also indicated trans-male nuclei carry less RNA, with fewer genes detected despite high sequencing saturation levels (Fig. 1G, S2B, S2C). Pathway analysis showed no indication for cell stress, apoptosis, or degradation, while instead revealing an upregulation of splicing factors, which was further corroborated by an increased rate of RNA-splicing based on reads mapping to exonic regions (Fig. 1H, S2C, D). Additionally, there was an overall downregulation of genes related to protein translation, which coincided with a higher degree of nuclear condensation, collectively indicating a decrease in transcriptional and translational output under androgen exposure (Fig. 1I, S2D).

### Hormone responsive luminal cells lose expression of the progesterone receptor and activate male-specific gene programs

To further understand the impact of testosterone treatment, we turned to luminal-HR^+^ cells, which were the only cell type to express all three hormone receptors (*AR, ESR1* and *PGR*), and which showed the largest treatment-induced change in chromatin and RNA (Fig. 1B, S2E, F). This subclass of KRT8^+^ luminal cells formed six subclusters based on RNA expression, with the cells from cis-female and trans-male samples distributing distinctly across these subgroupings (Fig. 2A, S4A). RNA-velocity analysis also suggests that cis-female and trans-male cells follow diverging trajectories and terminate at two distinct transcriptional states (Fig. 2A, B). The subcluster containing most trans-male cells (lup 2) was enriched for fatty acid metabolism and calcium signaling pathways, while the major cis-female subclusters (lup 1, lup 3) were associated with growth factor and estrogen signaling as well as mammary gland development (Fig. S4B). Of two minor subclusters, one showed expression of DNA replication and cell cycling genes (labeled: cycling), while the other was enriched for ribosomal genes (labeled: ribo) (Fig. S4B, Table 3). The largest subcluster shared by trans-male and cis-female cells (lup 4) showed a unique expression pattern marked by androgen response genes, steroid metabolism, and low *ESR1* expression (Fig. 2A, B, C, S4B). Various trajectory analyses implied lup 4 might present a less differentiated cell state from which the main cis-female and trans-male subclasses emerge (Fig. 2B, S4C, D). This is in accordance with reports that luminal progenitors are ER^-^ and that lipid metabolism is emerging as a key regulator of progenitor cell maintenance^3, 17^.

**Figure 2:**
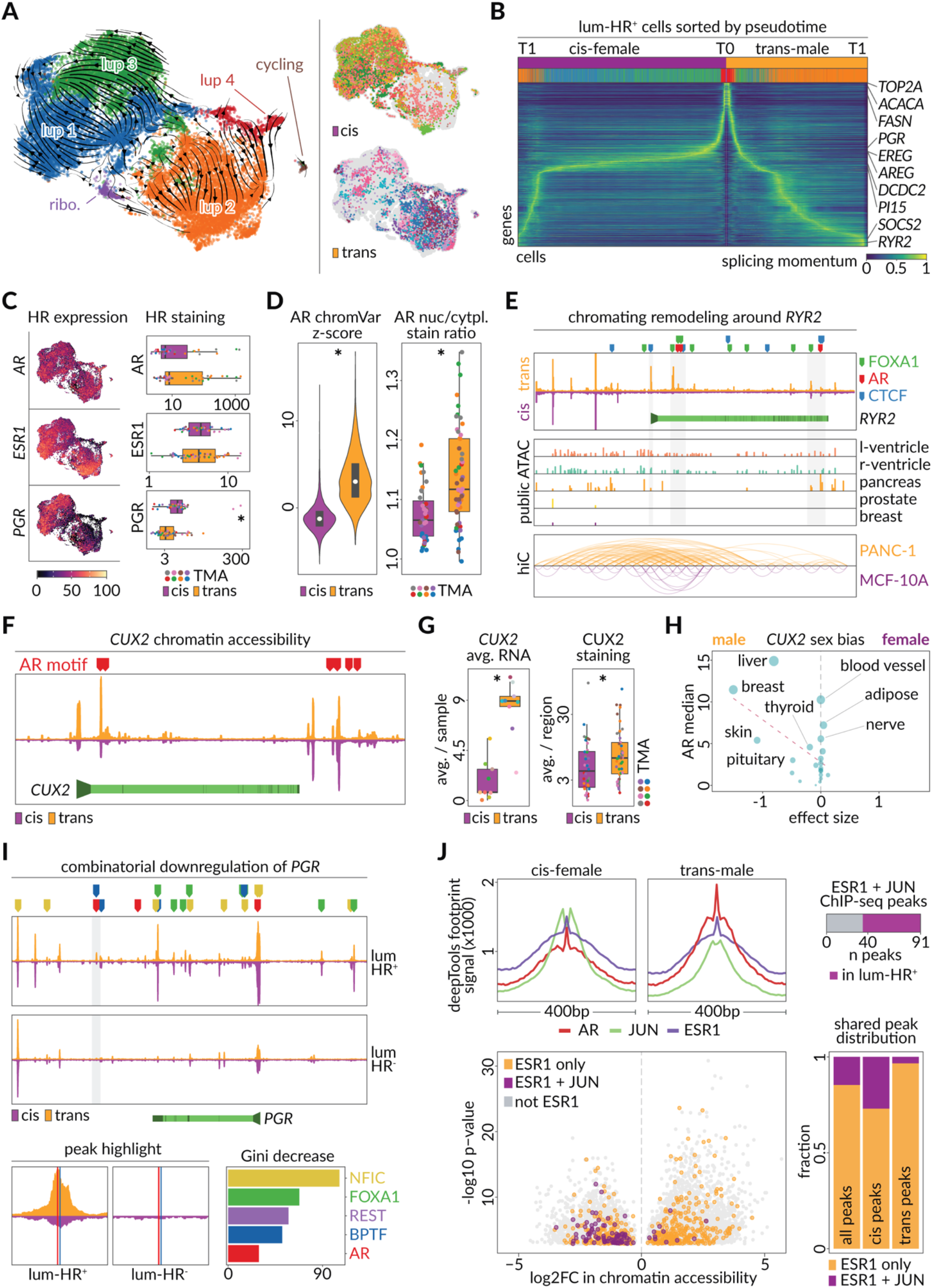
Hormone responsive epithelial cells of the trans-male breast are altered for genes that show sex bias in other tissues. **A)** snRNA-seq UMAP showing detected subclusters of hormone receptor expressing luminal cells (luminal-HR^+^), with RNA-velocity streams overlayed and gender identity shown by sample (right panels). **B)** RNA-velocity pseudotime ordering of trans-male and cis-female luminal-HR^+^ cells. Time 0 (T0) in the center and respective endpoints of cis-female and trans-male lineages (T1) at the outer maxima. Annotation bars show gender identity and subcluster assignment of each cell. Rows are annotated with highly differentially expressed genes or subcluster markers. **C)** Left panel overlays hormone receptor RNA expression on UMAP from (a). Right panel shows boxplots of hormone receptor staining intensities averaged across luminal-HR^+^ cells in CODEX microarray regions of cis-female (purple) and trans-male (orange) tissues. Dot colors indicate from which TMA the shown regions originate. (p-value, Wilcoxon: PGR = 0.00041) **D)** Violin plot (left) showing per cell chromVAR motif enrichment z-scores for AR (CisBP M03389_2.00) from luminal-HR^+^ cells in snATAC-seq (p-value, Wilcoxon < 2.2 x 10^-16^). Boxplot (right) showing average nuclear to cytoplasmic staining ratio for AR in luminal-HR^+^ cells from each TMA region (p-value, Wilcoxon = 0.00021). Dot colors indicate from which TMA the shown regions originate. Purple corresponds to cis-female and orange corresponds to trans-male data. **E)** *RYR2* chromatin accessibility (top) for cis-female (purple) and trans-male (orange) luminal-HR^+^ cells, with highlighted motif binding sites of AR, FOXA1 and CTCF. The *RYR2* gene body (light-green) is shown with promoter (arrow) and exon boundaries (dark-green). Also shown (center) is chromatin accessibility data for the same genomic region in tissues with varying *RYR2* expression (right ventricle, left ventricle and pancreas, prostate, breast) and Hi-C data (bottom) comparing three-dimensional chromatin structure of the same region in PANC-1 (pancreas) and MCF-10A (breast) cell lines. **F)** AR motif binding sites (red markers) across open chromatin regions around the *CUX2* locus of trans-male (orange) and cis-female (purple) luminal-HR^+^ cells. *CUX2* gene body (light-green), exon boundaries (dark-green) and promoter (arrow) are shown below. **G)** Per-sample average RNA (left, adjusted p-value, MAST < 2.2 x 10^-16^) and per-region average staining intensity (right, p-value, Wilcoxon = 0.027) of *CUX2* in cis-female (purple) and trans-male (orange) tissues. Dot colors indicate from which TMA the shown regions originate. **H)** Effect sizes of *CUX2* sex bias in GTEx tissues shown as a function of median *AR* expression (vertical axis and dot-size). Positive and negative values indicate female and male bias, respectively. **I)** Chromatin accessibility (top) around the *PGR* locus in trans-male (orange) and cis-female (purple) luminal-HR^+^ and luminal-HR^−^ cells. The *PGR* gene body (light-green) is shown with promoter (arrow) and exon boundaries (dark-green). Bottom left shows a magnified view of an AR motif containing peak (location highlighted in gray above). On the bottom right are the importance levels of transcription factors that co-bind with AR and determine the directionality of the transcriptional change in the target (as identified through fitting a random forest to classify upregulated or downregulated AR-bound genes using the accessibility of the peaks of each gene at the occurrences of motifs). Binding sites of these transcription factors are visualized with colored markers in the genomic region above. **J)** Top left panel shows footprint of AR (red), JUN (green), and ESR1 (purple) in cis-female (left) and trans-male (right) luminal-HR^+^ cells. Bottom left shows the average log2FC of chromatin accessibility in peaks containing only the ESR1 motif (orange), ESR1 and JUN motifs (purple), or no ESR1 motif (gray). Bottom right shows the fraction of JUN motif overlapping peaks among all luminal-HR^+^ peaks (left), cis-female–specific luminal-HR^+^ peaks (middle), or trans-male–specific luminal-HR^+^ peaks (right) containing both ESR1 and JUN motif (purple) or only ESR1 motif (orange). Top right shows the fraction of peaks with both ESR1 and JUN motifs which had *in vitro* ChIP-seq evidence for binding of both JUN and ESR1 (purple).

Throughout androgen treatment, *AR* and *ESR1* RNA levels remained stable, while *PGR* was downregulated both in RNA and protein levels (Fig. 2C, S4A, S2E). However, treatment induced AR activity was detected through chromatin accessibility around androgen response elements (AREs) and when comparing nuclear- to cytoplasmic AR signal intensity, which remained unaltered for ESR1 and PGR (Fig. 2D, S4E, F). Differentially accessible chromatin in trans-male samples further revealed strong enrichment of AREs alongside motifs of several members of the forkhead family of transcription factors (Fig. S4G). The AR pioneer factor, FOXA1, showed the strongest enrichment in trans-male specific chromatin, indicating it may aid AR-initiated changes including the strong upregulation seen in the Ryanodine receptor calcium channel *RYR2* gene (Fig. 2E, S3, S4H, I, Table 4)^18^. RYR2 regulates calcium signaling events in the heart, blood vessels and pancreas, and the *RYR2* loci in these organs show accessibility of AR and FOXA1 targeted regions that are natively inaccessible in breast tissue, indicating AR might induce molecular programs of other AR responsive tissues through chromatin remodeling (Fig. 2E, S5B)^19, 20^. We hypothesize that activated calcium signaling in Luminal-HR^+^ cells results in enhanced secretion of numerous ligands, relaying indirect effects of androgen therapy on other cell types through paracrine signaling (Fig. S4I, S5I and 6G).

AR activation also seemed to significantly alter chromatin accessibility around the most upregulated transcription factor *CUX2* (Fig. 2F, G, S3, Table 4). *CUX2* normally shows the highest expression in the cis-male–exclusive prostate tissue, but *CUX2* also exhibits higher expression in cis-male liver and cis-male breast samples from the Genotype-Tissue Expression (GTEx) database compared to corresponding cis-female tissues, indicating androgen activates cis-male-biased expression patterns (Fig. 2H, S5A). In fact, the top up- and down-regulated genes (|log2FC| > 0.5) in luminal-HR^+^ cells showed strong cis-male and -female sex bias, respectively, in GTEx data for breast and other organs as well (Fig. S5B). Further, many of these genes were found differentially regulated when comparing epithelial enriched cis-male and cis-female breast samples, corroborating the activation of cis-male- and suppression of cis-female-specific genes by androgen treatment (Fig. S6A-E). This analysis also found fewer genes were detected in cis-male GTEx samples, similar to our findings above showing reduced transcriptional output under androgen exposure (Fig. S2C, S6B).

Given the repression of female gene expression programs, we also examined how androgen interferes with the other sex hormone receptors, PGR and ESR1. The majority of trans-male patients experience cessation of menses within 6 months of treatment, so we first examined PGR activity, since this receptor is critical for ovulation and was the most down-regulated transcription factor in androgen-treated luminal-HR^+^ cells (Fig. S3, Table 4)^21, 22^. Chromatin data suggests AR directly binds the *PGR* promotor alongside other accessible regions, indicating a potential for direct repression, and a broader random forest analysis indicates the presence of likely co-repressors at such AR-bound downregulated genes (Fig. 2I, S5C). Of these, NFIC and FOXA1 show co-regulatory function in the prostate^23, 24^. Surprisingly, PGR motif accessibility increased following androgen treatment, which is likely an artifact of sequence similarity between the DNA binding domains of AR and PGR (Fig. S4G, S5E)^25^. Indeed, PGR binding sites were enriched near AREs, with only the glucocorticoid receptor (NR3C1) showing a stronger linkage, and a linear regression analysis confirmed PGR motif accessibility could not be predicted from its gene expression, unlike for the other nuclear receptors studied (Fig. S5D, E).

Despite being a highly expressed hormone receptor and an archetypical female gene regulator, ESR1 binding activity appeared unaltered by testosterone (Fig. 2J, S2E, Table 4). However, estrogen signaling pathways were downregulated, potentially because the AP-1 transcription factors that help carry out this signaling were reduced in activity, particularly BATF^26, 27^ (Fig. S4B, G, S3, S5F, Table 4). Furthermore, the archetypical AP-1 factor JUN, which physically interacts with both ESR1 and AR, showed significant decrease in both expression and motif accessibility (Fig. 2J, S3, Table 4)^28, 29^. We found that among the ESR1 peaks, those that have a match to the JUN sequence motif become less accessible upon androgen treatment. Most of these accessible chromatin peaks also have ChIP-seq evidence for binding of both ESR1 and JUN at the same genomic positions, further suggesting co-regulatory activity (Fig. 2J). Amongst the targets that are potentially downregulated through reduced ESR1 and AP-1 factor binding, we found the two important growth factors *AREG* and *EREG*, both of which are estrogen responsive in breast cancer (Fig. S5G, H, S3, S6E, Table 4)^30–36^. However, at the protein level AREG increased and became more concentrated in luminal-HR^+^ cells (Fig. S5I). Secretion of AREG depends on cleavage of membrane bound pro-AREG via matrix metalloproteases such as ADAM17, which was found downregulated in the neighboring luminal-HR^-^ and basal cells, potentially causing an accumulation of uncleaved pro-AREG in luminal-HR^+^ cells^37, 38^ (Fig. S5J). Indeed, this could also explain why the AREG receptor EGFR was downregulated in neighboring basal cells but upregulated in luminal-HR^+^ cells (Fig. S5H, K).

### Fibroblasts alter signaling to epithelial cells through reduced laminin production

We also studied fibroblasts, which are one of two other AR-expressing breast cell populations and can direct breast epithelial cell behavior through control of ECM composition (Fig. S2E)^39, 40^. Based on RNA expression, breast fibroblasts consist of two matrix subtypes, a previously described lipo-fibroblast group expressing *PPARG*, and a population we termed vascular-like that expresses genes such as *NRP1* (normally expressed on endothelial cells). Chondrocytes were also detected (Fig. S7A, B, Table 3)^41, 42^. Similar to luminal-HR^+^ cells, fibroblasts showed increased accessibility of AR-motif containing chromatin, centrality of AR motif occurrence with respect to peak summit, and increased ratios of AR nuclear to cytoplasmic staining levels (Fig. S7C, D, E). Staining also revealed increased proportions of an AR expressing (fibr-main) and an epithelial associated (fibr-epi) fibroblast subgroup that stained for keratin 8 and 23 due to epithelial cell proximity (Fig. S7F, G).

The ECM constituent laminins *LAMB1* and *LAMA2* were among the most downregulated genes in fibroblasts, which was consistent with differences between fibroblast enriched GTEx cis-male and cis-female breast samples (Fig. S3D, S6E, S7H, Table 4). The genomic loci of these laminins revealed increased ARE accessibility, implying direct AR mediated repression (Fig. S7I). Indeed, LAMB1 staining was also reduced in LAMB1^+^ fibroblasts, while LAMA2 protein levels were unaltered (Fig. S7F, J, K). Interestingly, we found the *ITGB1* integrin receptor for LAMB1 and LAMA2 was specifically downregulated in basal epithelial cells, which directly interface with the ECM and appear surrounded by a layer of LAMB1 (Fig. S7K, L)^43, 44^.

### The breast epithelium shows reduced smooth muscle coverage and becomes infiltrated by various stromal cell types in response to androgen

Given the observed changes in ECM composition and its structural relationship with the epithelium, we also explored how non-hormone sensing basal and luminal cells respond to androgen treatment (Fig 1B, S1H, S8A). While their relative proportions and primary spatial arrangement appeared unchanged, we noticed morphological alterations in acini and ductal structures (Fig 3A, S8B). Overall, epithelial cells of transgender male patients presented with enlarged nuclei and image analysis revealed corresponding acini were significantly smaller after androgen treatment (Fig. 3B, C, S8C). This also confirmed the typical smooth muscle layer surrounding these structures was reduced in transgender men, which may be linked to the above-mentioned loss of AREG as its loss is reported to cause reduced myoepithelial coverage in the mammary duct (Fig. 3A, C, S8C)^45^.

**Figure 3:**
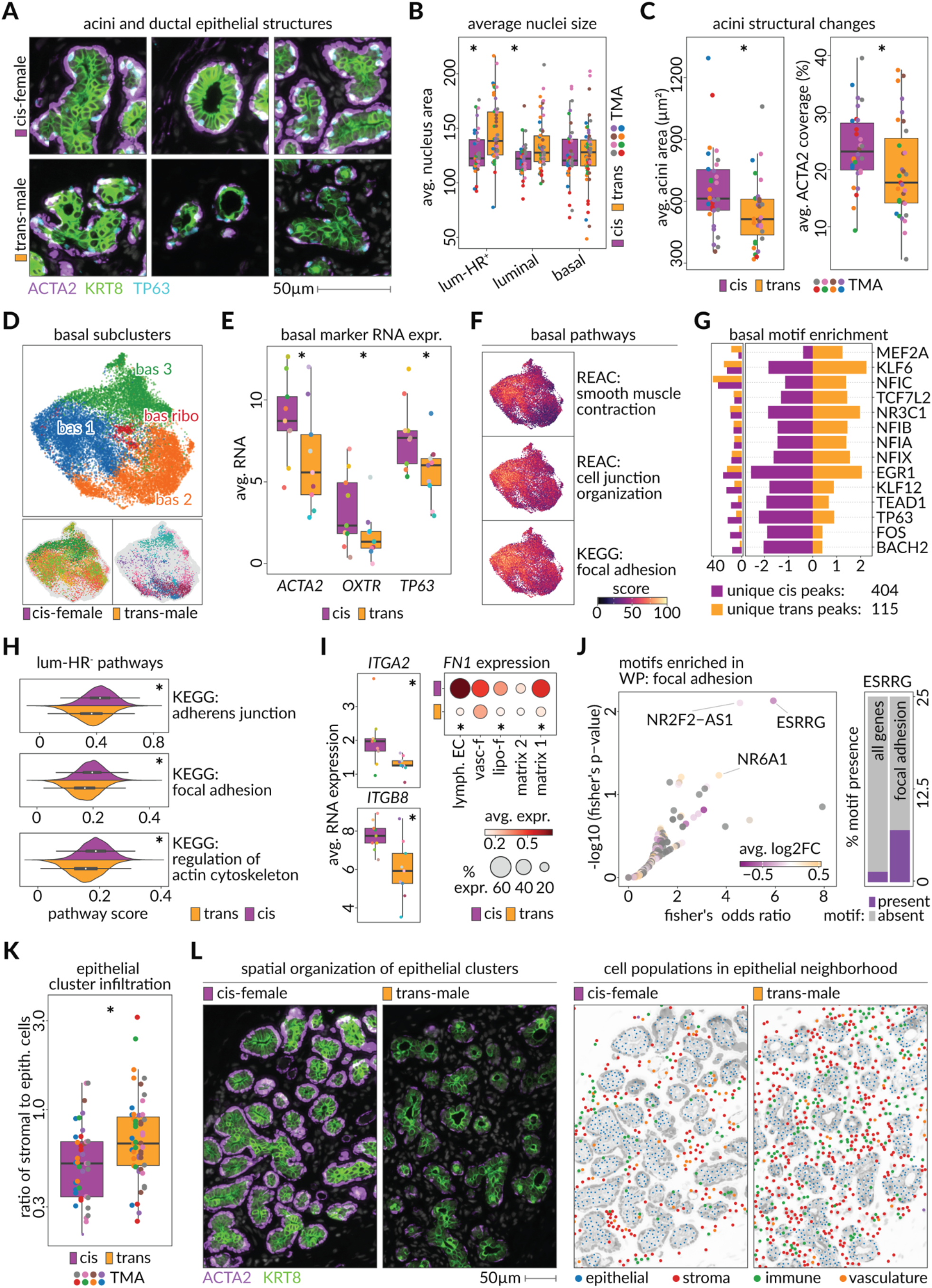
Epithelial cells without hormone responsiveness loose contractile functions upon hormone replacement therapy. **A)** Images from CODEX microarray data showing mammary acini structures from cis-female (top) and trans-male (bottom) tissues marked by expression of ACTA2 (basal cells; purple) TP63 (basal cell nuclei; blue) and KRT8 (luminal cells; green). **B)** Boxplots show per region average nucleus area among cis-female (purple) and trans-male (orange) epithelial cells (luminal-HR+, luminal cells, and basal) (p-values, Wilcoxon: luminal-HR+ = 0.00064, luminal = 0.016, basal = 0.54). Dot colors indicate TMA origin of shown regions. **C)** Boxplots show average area of acinar structures (left panel) and average area of acini border that was filled with ACTA2 signal (see Fig. S8C) among cis-female (purple) and trans-male (orange) tissues of CODEX microarray data. Dot colors indicate TMA origin of shown regions. (p-values, Wilcoxon: area = 0.026, ACTA2 coverage = 0.012) **D)** UMAP showing subclusters of basal cells in snRNA-seq data (top) and their distribution across trans-male and cis-female samples (bottom). **E)** RNA expression of ACTA2, OXTR (lactation markers) as well as TP63 in basal cells of trans-male (orange) and cis-female (purple) samples (adjusted p-values, MAST: ACTA2 = 8.86 x 10-296, OXTR = 9.59 x 10-262, TP63 = 1.16 x 10-96). **F)** Module scores of significantly enriched pathways overlaid over the UMAP of basal cells (REAC = Reactome, KEGG = Kyoto encyclopedia of genes and genomes). **G)** Right panel shows the enrichment of motifs among unique accessible chromatin peaks of trans-male (orange) and cis-female (purple) basal cells. Left panel shows the fraction of the peaks of the corresponding cells which overlap with the motif. **H)** Kernel density estimation and boxplot of module scores for selected significantly altered structural pathways in luminal-HR– cells (p-values, Wilcoxon: KEGG: adherens junction = 4.13 x 10-285, KEGG: focal adhesion = 1.42 x 10-255, KEGG: regulation of actin cytoskeleton < 1.42 x 10-255). **I)** Boxplots show average RNA expression (top) of integrin receptors from the “KEGG: regulation of actin cytoskeleton” pathway in luminal-HR– cells (adjusted p-values, MAST: ITGA2 = 4.89 x 10-201, ITGB8 = 6.40 x 10-267) and average expression of the ITGA2 and ITGB8 ligand FN1 in fibroblast subclusters and lymphatic endothelial cells (bottom) of trans-male and cis-female samples (adjusted p-values, MAST: matrix 1 = 1.66 x 10-54, matrix 2 = n.s., lipo-f = 1.32 x 10-16, vasc-f = n.s., lymph. EC = 3.13 x 10-99). **J)** Scatterplot shows Fisher’s exact test odds ratio (x-axis) and –log10 p-value (y-axis) corresponding to enrichment of each motif among the chromatin accessibility peaks for the genes of the “WikiPathways: focal adhesion pathway”. Colors indicate the log2 fold change in gene expression of the transcription factor corresponding to each motif. Gray motifs correspond to transcription factors without differential gene expression among luminal-HR– cells. Right barplot shows the fraction of all genes (left) and genes annotated within the focal adhesion pathway (right) which contain a chromatin accessibility peak matching the ESRRG sequence motif (cisBP ESRRG_697). **K)** Boxplot shows the ratio of stromal to epithelial cells in the epithelial neighborhood (see. Fig. S8C) among regions of cis-female (purple) and transgender male (orange) tissue in CODEX microarray data. (p-value, Wilcoxon = 0.0052). Dot colors indicate TMA origin of shown regions. **L)** The left microscopic image shows luminal (green KRT8-stained) and basal (purple ACTA2-stained) cells in a cis-female (left) and trans-male (right) breast tissue. The right schematic indicates the epithelial (blue), stromal (red), immune (green), and vasculature (orange) components of the left image.

As a whole, basal/myoepithelial cell proportions were unchanged, but androgen treatments did transition cells away from two subclusters that were cis-female dominated (bas 1 & bas 3) and towards one enriched for trans-male samples (bas 2), with the only other cluster expressing ribosomal genes (bas ribo) similarly to luminal-HR^+^ (Fig. 3D, Table 3). Particularly, basal cell expression of the lactation-associated genes *ACTA2* and *OXTR* as well as *TP63* were all reduced at the RNA and protein level suggesting contractile impairment (Fig. 3E, S8D)^46–48^. *OXTR* was also found reduced in epithelial enriched cis-male breast GTEx samples suggesting these changes result from androgen signaling (Fig. S6). RNA velocity analysis further indicates that androgen-treated cells follow a unique developmental trajectory and converge at a distinct terminal state where a reduction of genes relating to smooth muscle contraction, focal adhesion and cell junction organization is seen (Fig. 3F, S8E). For the latter pathway, this likely results from lowered chromatin activity and gene expression of the BACH2 transcription factor, which is enriched for binding sites at corresponding gene loci and which is also lowered in GTEx cis-male breast tissue (Fig. 3G, S6, S8F, G, Table 5).

Among the three epithelial cell types, the luminal-HR^-^ population showed the lowest margin of differential expression and none of the detected subclusters showed notable population bias (Fig. S3, S8H). Like in the luminal-HR^+^ population, we detected a subcluster with expression of cell cycling genes (labeled: lun cycling) and another group with high expression of ribosomal genes (labeled: lun ribo), while the remaining clusters were distinguishable by various growth-factor and immune related pathways (Fig. S8H, I, Table 3, Table 4).

Similar to the basal population, structural pathways of focal adhesion, adherens junctions and actin cytoskeleton regulation were downregulated by androgen (Fig. 3H). As described above, many structural pathways are modulated through integrin transmitted instructions from the ECM, and significant loss of the candidates *ITGA2* and *ITGB8* in trans-male luminal-HR^-^ cells and in epithelial enriched cis-male breast samples were observed (Fig. 3I, S6). When matching ligands secreted in other cell types, we found decreased expression of the ECM component and critical cell adhesion and morphology regulator *FN1* in subtypes of fibroblasts and also in lymphatic endothelial cells (Fig. 3I)^49–51^. While staining intensity of FN1 was not decreased in these cell types after androgen treatment, there were fewer FN1^+^ fibroblasts identified in the immediate epithelial neighborhood (Fig. S7F, S9A, B). When assessing which transcription factors might be involved, we found motif over-representation of Estrogen Related Receptor Gamma (ESRRG) binding motifs in the chromatin peaks of focal adhesion genes, which is downregulated in trans-male luminal-HR^-^ cells (Fig. 3J, S9C). ESRRG is closely related to ESR1 and can be suppressed by androgen in AR^+^ and AR^-^ prostate cancer cells^52, 53^.

### Increased thrombospondin 1 production by adipocytes and fibroblasts correlates with decreased capillary vasculature during androgen exposure

While studying the epithelium, we observed trans-male samples harbor a significantly higher amount of non-epithelial cells proximal to acini and ducts, leading to decreased epithelial cell density in these structures (Fig. 3K, L). With fibroblasts being the largest group found in the inter-epithelial spaces, this observed infiltration might explain previously reported fibrosis in transgender male breast tissue (Fig. S9C)^54, 55^. Other notable populations found in the epithelial neighborhood were immune and endothelial cells, with the latter group showing a noticeable decrease in presence (Fig. S9C, Fig. 4A).

**Figure 4:**
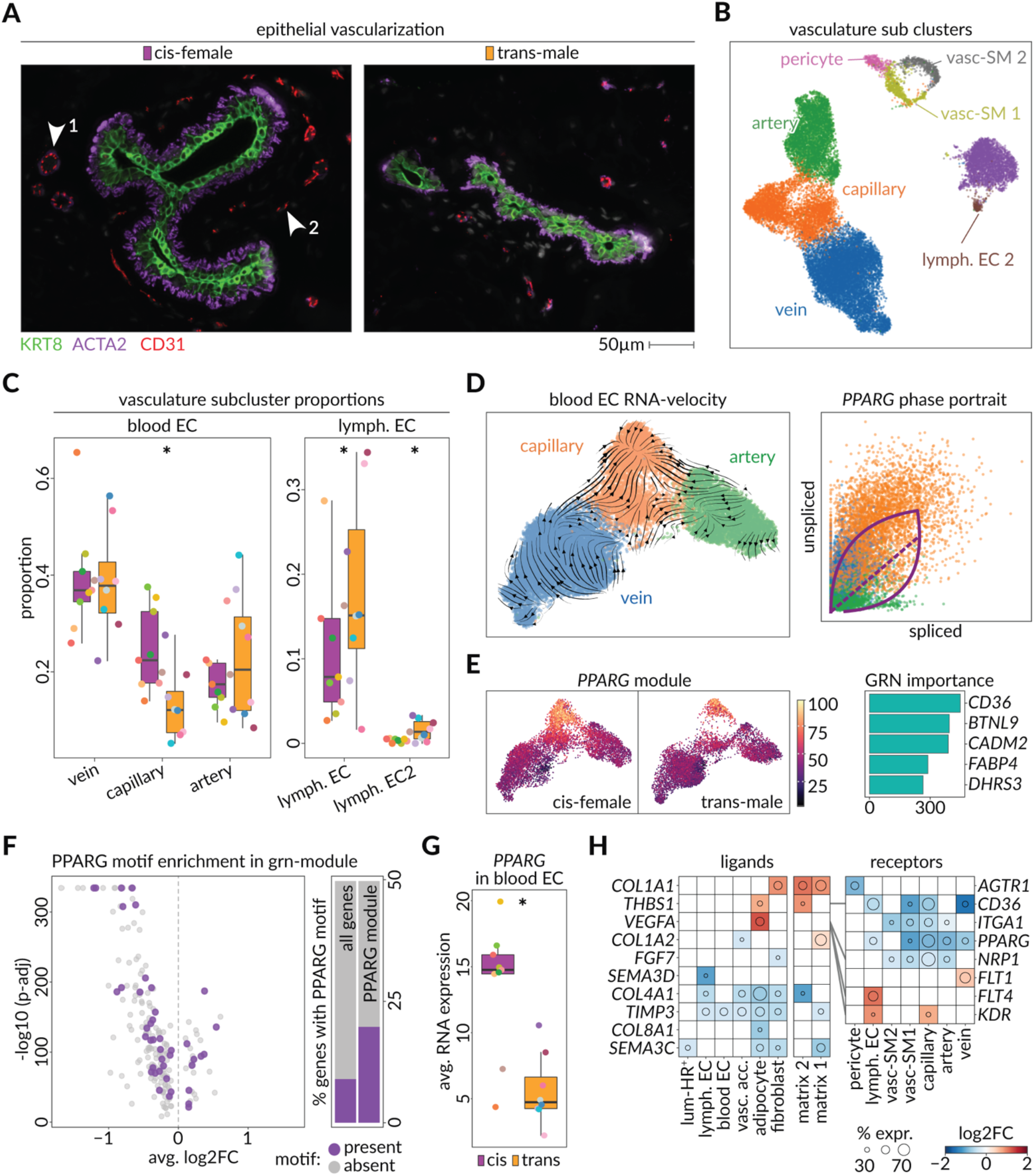
Hormone replacement therapy reduces epithelial vascularization through PPARG activity. **A)** Microscopic images show vascularization of two ductal structures in a cis-female (left) and a trans-male (right) breast tissue in CODEX microarray data. KRT8 (green) marks luminal cells, ACTA2 (purple) marks green cells, and CD31 (red) marks endothelial cells. Arrows point out 1: larger vessels with smooth muscle layer and 2: smaller vessels without smooth muscle layer. **B)** UMAP shows vasculature subclusters detected in the snRNA-seq dataset. (left = blood endothelial cells, right = lymphatic endothelial cells, upper-mid = vascular accessory cells). **C)** Boxplots show the proportions of vascular subclusters in each sample of the snRNA-seq data, split by gender-ID (GLM p-values, generalized linear model fitting a poisson: vein = 6.72 x 10^-45^, capillary = 5.31 x 10^-77^, artery = 3.58 x 10^-5^, lymph. EC = 1.33 x 10^-22^, and lymph. EC 2 = 0.0071) **D)** UMAP (left) shows blood endothelial cells overlaid with scvelo stream plots. The scatterplot shows the ratio of spliced (horizontal axis) and unspliced RNA molecules (vertical axis) of *PPARG* among vein (blue), capillary (orange), and artery (green) blood endothelial cells. Dashed diagonal indicates the steady state ratio. Top and bottom arcs indicate the estimated kinetic parameters of *PPARG* induction and repression, respectively. **E)** *PPARG* gene regulatory network module score overlaid on UMAP plot among cis-female (left) and trans-male (right) blood endothelial cells. Barplot shows GRN importance scores of the top 5 genes coexpressed with *PPARG*. **F)** Volcano plot shows the average log_2_ fold change and –log_10_ adjusted p-value for differential expression of genes within the *PPARG* module among the trans-male and cis-female blood endothelial cells. Purple data points indicate genes with a chromatin accessibility peak overlapping the PPARG transcription factor sequence motif (CisBP PPARG_676) match. Barplots show the fraction of all genes (left) or genes within *PPARG* module (right) which contain a chromatin accessibility peak overlapping the PPARG transcription factor sequence motif (purple). **G)** Boxplot shows average expression of *PPARG* in blood endothelial cells of cis-female (purple) and trans-male (orange) samples in snRNA-seq data. **H)** Heatmap shows the log_2_ fold change in expression of ligand (left) - receptor (right) pairs among cell types and vascular subclusters the trans-male and cis-female samples. Colors indicate log2 fold change in expression, and diameter of the circle shows the percent of cis-female cells expressing the gene.

Transcriptomic data suggests the breast vasculature is characterized by three distinct cell types. Blood endothelial cells (blood EC) consisting of arterial, capillary, and venous cells, were called according to previously defined signatures^56^. Lymphatic endothelial cells expressing *PDPN, LYVE1* and *FLT4* were separated into two distinct subclusters (lymph. EC & lymph. EC2), and pericytes grouped together with two subclusters of vascular smooth muscle cells (vasc. SM1 & vasc. SM2) (Fig. 4B, S10A, B, C). Through antibody staining we found the general CD31^+^ endothelial population further separated into an ACTA2^+^ contractile subgroup (endo-SMA), a CD45^+^ immune associated subgroup (endo-immu.) and another group showing high levels of LNX1 and CD36, resembling the expression of capillaries (endo-LNX1^+^). Lymphatic ECs were classified via LYVE1 and PDPN (Fig. S10D, E).

While lymphatic endothelial cells were only marginally represented in the epithelial neighborhood, we found their overall proportions within the vasculature were increased based on RNA, ATAC and tissue staining data (Fig 4C, S10F, G, H). Thus, blood endothelial cells were the main constituent of epithelial-adjacent vascularization, and we found that the proportions of arteries and veins remained stable during testosterone treatment (Fig. S9C, 4C). However, trans-male vasculature was clearly reduced for capillaries, which was corroborated by the specific reduction of the LNX1^+^ endothelial subclass in the epithelial neighborhood (Fig.4A, C, S10F, H).

To understand mechanistically why capillary cells were reduced by testosterone, we performed RNA-velocity analysis on blood-ECs. and found capillaries harbor a distinct terminal state of differentiation that appears driven by PPARG (Fig. 4D). This relationship was further corroborated by the specific expression of an inferred PPARG gene regulatory network (GRN) in capillaries, which contained PPARG-associated lipid-metabolism markers *FABP4* and *CD36* and was enriched for blood vessel development and angiogenesis pathways, all of which showed enrichment for PPARG motifs near their associated loci (Fig 4E, F, S10I)^57, 58^. Together with decreased expression of *PPARG* and reduced activity of the *PPARG* GRN-module, this implies that *PPARG* loss drives the reduced angiogenesis and capillary proportions that result from androgen exposure (Fig. 4F, G). To study external factors that might promote the initiation of this regulatory mechanism, we selected highly expressed receptors for the different vascular subclusters and matched them to differentially expressed ligands from all remaining breast cell types. This found that androgen-sensitive adipocytes and matrix 2-type fibroblasts upregulate the antiangiogenic CD36 ligand Thrombospondin 1 (*THBS1*), which is also increased in adipose and fibroblast enriched GTEx cis-male breast samples (Fig. 4H, S6)^59, 60^.

A similar ligand receptor analysis to that above found the vascular growth factor *VEGFA* is also upregulated in adipocytes in response to androgen, and that lymphatic endothelial cells were the only cell types with increased expression of both the corresponding receptor *KDR* (VEGFR2) and *FLT4*, which dimerize to promote angiogenesis (Fig. 4H)^61, 62^. These results suggest this mechanism underlies the increased lymphatic endothelial cell numbers observed, which is supported through pathway analysis that found VEGFR2 mediated cell proliferation is higher in these cells (Fig. S10J).

### Androgen treatment reduces macrophage numbers in the transgender male breast, while T-Cells become more associated with epithelial structures

In addition to fibroblast and endothelial cell compartments, immune cells were also found well represented near the breast epithelium (Fig. S9C). Sub-clustering of immune populations represented in the nuclei data revealed the lymphoid compartment consisted of CD8^+^ and CD4^+^ T-Cells, T-effector cells, NK cells and two classes of B-Cells, while the myeloid compartment consisted of monocyte derived dendritic cells (labeled: mono.DC), macrophages, dendritic cells (labeled: DC) and monocytes. A small cluster of HSCs was also detected (Fig. 5A, B, S11A, B).

**Figure 5:**
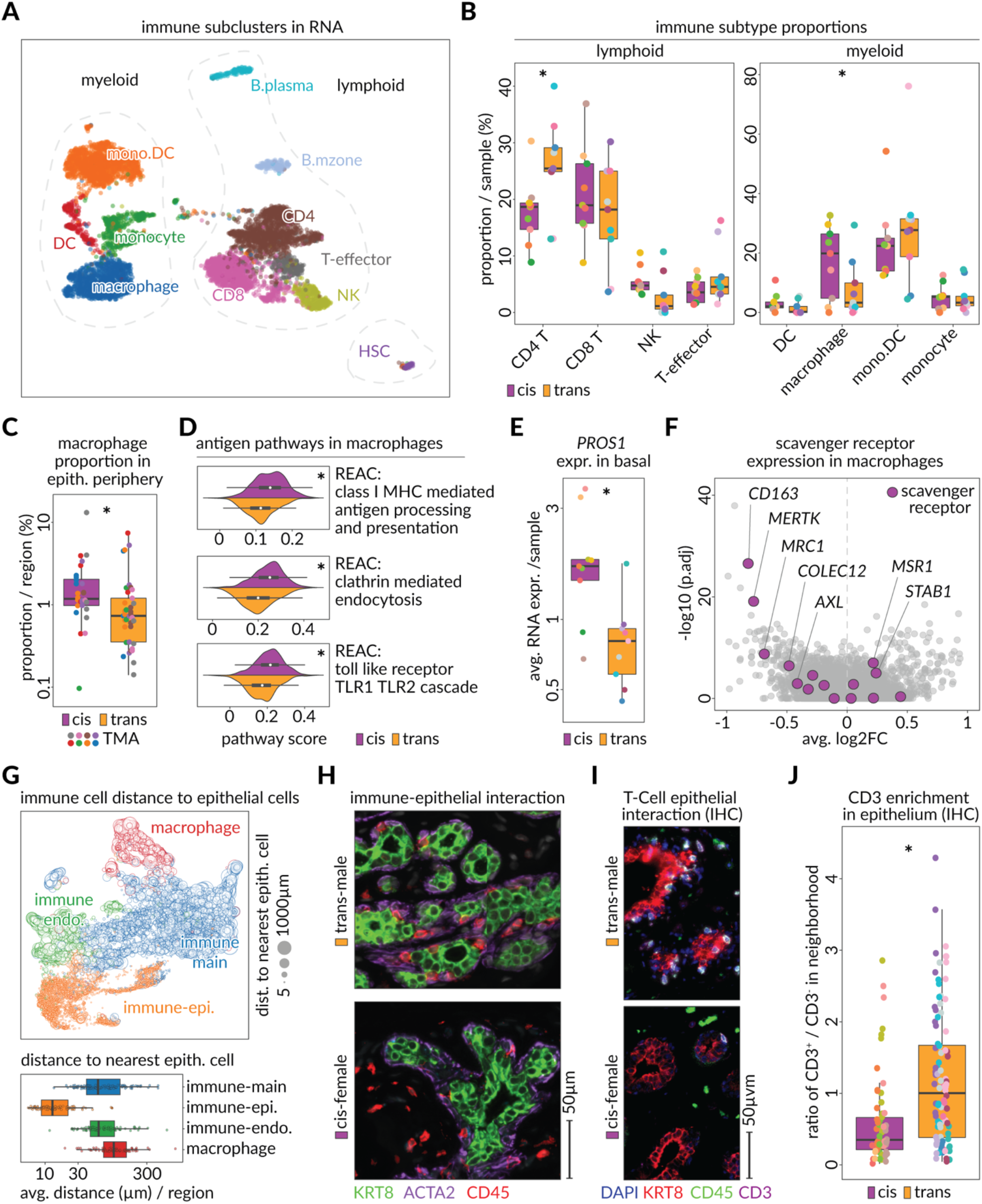
Hormone replacement therapy dominates helper T lymphocytes and reduces the presence of innate immunity. **A)** UMAP showing subclusters of all myeloid (left) and lymphoid (right) cells detected in the snRNA-seq data. (CD8 = CD8^+^ T-Cells, CD4 = CD4^+^ T-Cells, T-effector = Effector T-Cells, NK = natural killer cells, mono.DC = monocyte derived dendritic cells, DC = dendritic cells) **B)** Boxplots show the fraction of main immune cell subtypes within entire immune compartment in each sample (GLM p-values, generalized linear model fitting a poisson: CD4 = 0.00035, CD8 = 4.035 x 10^-13^, T-effector = 0.045, NK = 0.00035, mono.DC = 0.017, macrophage = 0.52, monocyte = 0.055, DC = 0.0001). **C)** Boxplot shows the proportion of macrophages within the periphery of epithelial cells in cis-female (purple) and trans-male (orange) tissue regions of the CODEX microarray data. (p-value, Wilcoxon = 0.003) **D)** Kernel density estimates, and boxplots show the module scores of immune relevant Reactome pathways in macrophages of trans-male (orange) and cis-female (purple) samples. (p-values, Wilcoxon, class-I MHC mediated antigen processing/presentation = 8.32 x 10^-17^, clathrin mediated endocytosis = 3.64 x 10^-21^, roll like receptor TLR1 TLR2 cascade = 3.89 x 10^-16^) **E)** Boxplots show the average RNA expression of *PROS1* in basal cells of cis-female (purple) and trans-male (orange) samples (adjusted p-value, MAST = 3.17 x 10^-192^). **F)** Volcano plot shows the average log_2_ fold change and –log_10_ adjusted p-value assessing the differential expression of genes in trans-male macrophages compared to cis-female macrophages. Purple data points indicate scavenger receptors. **G)** UMAP shows four immune cell staining sub classes (macrophage; red, immune endo.: green, immune-main: blue, and immune-epi.: orange) according to the staining pattern in CODEX microarray data. Size of the data points indicates the distance to the most proximal epithelial cell. Boxplot (below) summarizes the average distance of each group of immune cells to their most proximal epithelial cell. **H)** Microscopic images show staining of luminal (KRT8; green), basal (ACTA2; purple) and immune cells (CD45; red) within a trans-male (top) and a cis-female (bottom) breast tissue in CODEX microarray data. **I)** Microscopic image shows IHC staining of luminal (KRT8; red), immune (CD45; green), and T-lymphocyte (CD3; purple) cells within a trans-male (top) and a cis-female (bottom) breast tissue. White cells are double positive for CD45 and CD3. **J)** Boxplot shows the ratio of immune cells (CD45^+^) expressing CD3 to those not expressing CD3 (T-lymphocytes vs. other immune cells) within the epithelial neighborhood of cis-female (purple) and trans-male (orange) breast tissues of IHC scan regions.

Overall, there were fewer macrophages amongst immune cells from transgender men, and they were specifically reduced in the epithelial neighborhood (Fig. 5B, C, S11C). Macrophages assist mammary tissue remodeling during ductal morphogenesis, and differential pathway analysis showed genes for antigen recognition, antigen presentation and endocytosis were expressed at lower levels in trans-male macrophages (Fig. 5D)^63, 64^. When examining ligand receptor pairs that might moderate these functions, we found basal cells strongly downregulated *PROS1*, which has been shown to stimulate inflammatory resolving efferocytosis in (Fig. 5E)^65^. The documented PROS1 receptors *MERTK* and *AXL* were also downregulated in macrophages, alongside multiple other scavenger receptors including *CD163, MRC1* and *COLEC12* (Fig. 5F)^66–68^. Notably, we also observed downregulation of *PROS1* in epithelial-enriched GTEx cis-male breast samples, providing further evidence of androgen-induced mediation of macrophage function (Fig. S6). Considering the reduction of mammary growth factors like AREG and EREG, these results suggest macrophages are less involved in clearance of regenerating epithelial tissue during androgen treatment.

In studying immune populations in the CODEX data, we found CD45^+^ cells also formed distinct subsets based on their adjacency to other cell types (Fig. 5G, S11D). Indeed, an epithelium-associated immune population found near or in direct contact with ducts and acini was identified, which was increased in proportion in trans-male tissues, and further analysis revealed this group was mainly constituted by T-Cells (Fig. 5G, H, I, J, S11E, F). Among T-Cell subsets found in our RNA data, the proportion of CD4^+^ T-Cells specifically increased, which was potentially driven by the upregulation of the T-Cell differentiation factor TCF7 (Fig. 5B, S11G)^69–71^. When searching for potential chemo attractants, we found that matrix-type fibroblasts upregulated *IL16*, which triggers a migratory response in CD4^+^ T-Cells (Fig. S11G)^72, 73^. Pathway enrichment of genes upregulated in CD4^+^ T-Cells further indicated that these cells are activated and proliferative in transgender men (Fig. S11H). While the role of CD4^+^ T-Cells in the normal breast remains largely unknown, these results might indicate their involvement in mediating the mammary epithelial immune microenvironment.

### Androgen treatment elicits lipid reduction through epithelial AZGP1 secretion and by directly targeting adipogenesis pathways in adipocytes

Although milk production relies mainly on cell types analyzed above, which constitute and surround acini and ducts, the human breast also contains significant amounts of adipose tissue (7-56%) that is found in large distinctly organized homogenous macro structures (Fig. S12A)^74^. To ensure inclusion of diverse cell types, adipose tissue was broadly removed during sample collection and adipocyte hydrophobicity resulted in their further exclusion during nuclei library preparations. Regardless, the captured adipose nuclei express the androgen receptor as expected and are thus of interest, especially considering the documented impact of androgens on energy metabolism and the fact that transgender men experience weight loss and redistribution of body fat (Fig. S2E, S12B)^13, 75–78^.

To explore how adipocytes might interface with their environment during androgen treatment, we performed a global ligand-receptor analysis, which found ligands and receptors belonging to the PI3K pathway are amongst the most frequently altered signaling components in our dataset (Fig. S12C). Across receptors of the PI3K pathway, the broadest upregulation was for the insulin receptor (*INSR*), as validated based on staining, with adipocytes showing the most significant change and the highest baseline expression of *INSR*. These results align with previous reports of increased insulin sensitivity and altered metabolism in transgender men (Fig. 6A, S12D, E)^13, 76, 79^.

**Figure 6:**
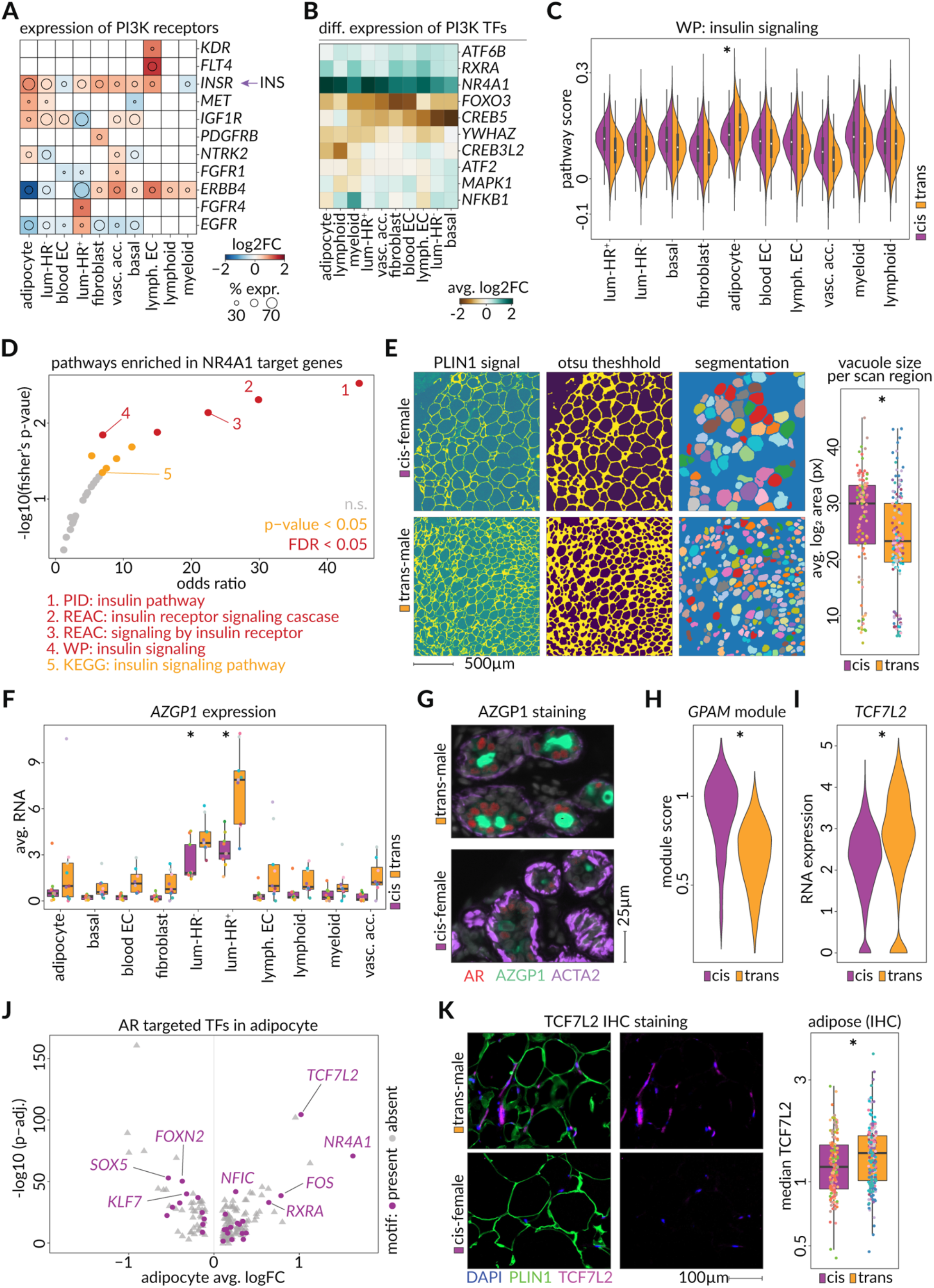
Testosterone induces PI3K pathway alterations with adipocytes showing distinct metabolic adaptations. **A)** Heatmap showing the log_2_ fold change in RNA expression of PI3K receptors (taken from “KEGG: PI3K-Akt signaling pathway”) among the 10 cell types of the breast. The diameter of the circle indicates the fraction of cis-female cells of the cell type expressing the receptor. INS indicates circulating insulin secreted in the pancreas. **B)** Heatmap showing the log_2_ fold change in RNA expression of the downstream transcription factors of the “KEGG: PI3K-Akt signaling pathway” among the 10 cell types of the breast. **C)** Pathway enrichment of tissue wide upregulated genes (> 7 cell types) that also have an NR4A1 motif in an enhancer (> 4 cell types). Horizontal axis shows the odds ratio (Fisher’s exact test) comparing frequency of selected genes in the pathway versus background and vertical axis shows –log_10_ p-value of Fisher’s exact test. FDR < 0.05 = red, p-value < 0.05 = yellow, n.s. = gray. (PID = Pathway Interaction Database, REAC = Reactome, WP = WikiPathways, KEGG: Kyoto Encyclopedia of Genes and Genomes). **D)** Module scores for the WikiPathways “WP: insulin signaling pathway” in all cell types, split by cis-female (purple) and trans-male (orange). (p-value in adipocytes, Wilcoxon = 7.59 x 10^-4^). **E)** Representative images of computational segmentation of lipid vacuoles (left, see methods), and resulting average area of adipocyte vacuoles per IHC scan region (p-value Wilcoxon = 0.00059). **F)** Boxplot shows the sample averages of *AZGP1* RNA-expression in each cell type in trans-male (orange) and cis-female (purple) samples (adjusted p-values, MAST: adipocyte = 5.95 x 10^-12^, basal = 2.42 x 10^-70^, blood EC = 8.67 x 10^-83^, fibroblast = n.s., luminal-HR^-^ = 4.54 x 10^-302^, luminal-HR^+^ = 0.00, lymph. EC = 2.83 x 10^-15^, lymphoid cells = 5.46 x 10^-12^, myeloid cells = 9.23 x 10^-8^). **G)** Microscopic image of a duct stained against AR (red), AZGP1 (green), and ACTA2 (purple) in a trans-male (top) and cis-female (bottom) breast tissue of the CODEX microarray. **H)** Violin plot shows the *GPAM* co-expression module (GRNboost2, 95^th^ percentile, p-value, Wilcoxon < 2.22 x 10^-16^) score in cis-female (purple) and trans-male (orange) adipocytes. **I)** Violin plot shows the *TCF7L2* expression in trans-male (orange) and cis-female (purple) adipocytes (adjusted p-value, MAST = 3.48 x 10^-105^) **J)** Volcano plot shows the differential expression of transcription factors in comparison of trans-male to cis-female adipocytes. Horizontal axis shows log_2_ fold change in expression and the vertical axis shows –log_10_ adjusted p-value. Purple data points indicate transcription factors with accessible chromatin matching the AR sequence motif (CisBP AR_689). **K)** Microscopic immunohistochemistry images show the staining against nuclei (DAPI; blue), adipocytes (PLIN1; green), and TCF7L2 (purple). Boxplot shows the median staining intensity of TCF7L2 among IHC scan-regions of cis-female (purple) and trans-male (orange) adipocytes (p-value, Wilcoxon = 0.0069).

To identify potential upstream regulators of insulin signaling, we investigated the PI3K related transcription factors and found that multiple candidates showed increased expression in multiple cell types (Fig. 6B). In particular, the orphan nuclear receptor *NR4A1* and its binding partner *RXRA* showed consistent upregulation, while the homeostatic regulator and target for inactivation by AKT, *FOXO3* and its nuclear chaperone *YWHAZ* were downregulated (Fig. 6B). *FOXO3* also showed lowered expression in GTEx cis-male breast samples (Fig. S6)^80–84^. Adipocytes had the highest baseline insulin signaling pathway activity score and were the only cell type showing a transcriptionally inferred increase in insulin signaling, further illustrated by upregulation of the critical PI3K components *PTEN*, *PIK3R1* and *AKT3* (Fig. 6C, S12F). Amongst PI3K related transcription factors, NR4A1 showed a particularly strong increase in adipocytes based on RNA and protein levels, and it was also upregulated in adipose-enriched GTEx cis-male breast samples (Fig. 6B, S6, S12G). NR4A1 has recently emerged as a novel target in multiple metabolic processes and cardiovascular function, and we found NR4A1 motif enrichment at genes associated with insulin signaling, supporting the notion that it might be an effector of glucose and lipid metabolism (Fig. 6D)^85–87^.

Insulin sensitivity is inversely correlated with adipocyte size, with hypertrophic adipocytes being indicative of metabolic dysfunction and insulin resistance, and our analysis revealed that androgen treatment does indeed lead to smaller lipid vacuoles (Fig. 6E)^88–90^. Considering this, lipolysis and lipogenesis related genes were analyzed, but these failed to stratify breast adipocytes according to their treatment status (not shown). However, an upregulation of the lipolysis regulator zinc-alpha glycoprotein 1 (*AZGP1*) was observed (Fig. 6F, G). AZGP1 is an androgen responsive secreted factor that is known for potently inducing lipid degradation and fat loss in smokers and cancer patients^91–94^. In trans-male samples, *AZGP1* expression was particularly increased in luminal cells, and it was also found increased in epithelial enriched GTEx cis-male breast samples (Fig. 6F, G, S6, S12H). Indeed, AZGP1 staining was also particularly visible in the cytoplasm of luminal-HR^+^ cells, indicating androgen treatment triggers fat reducing signals through AR responsive cells (Fig. 6G, S12H).

Adipocyte intrinsic factors potentially contributing to fat loss were also studied using gene regulatory networks, which found testosterone induces a downregulation of genes co-expressed with the metabolic marker *GPAM* (Fig. 6H). Pathway analysis of this module revealed an enrichment of genes involved in adipogenesis and adipocyte differentiation, implying adipocyte regeneration is constrained during androgen treatment (Fig. S12I). In searching for *GPAM*-module regulators we found accessible binding elements of the upregulated Wnt/B-catenin effector TCF7L2 were enriched in loci of the GPAM-module constituents (Fig. 6I, S12J). Besides being a repressor of adipogenesis, TCF7L2 is also crucial for glucose tolerance and insulin sensitivity^95–97^. In trans-male adipocytes, we found *TCF7L2* is a direct AR target and, alongside *NR4A1*, is one of the highest upregulated transcription factors in adipocytes (Fig. 6J). Adipose specific IHC staining also showed that TCF7L2 signal was increased in adipocytes of trans-male samples, further indicating some fat loss in transgender men may be due to an anti-adipogenic effect of androgen (Fig. 6K).

## Discussion

To understand the role that androgen plays in maintaining breast tissue homeostasis at the molecular level, we have performed a multimodal single-cell resolution profiling of breasts from transgender men undergoing gender affirming testosterone therapy. As our control samples were obtained from a combination of pre- and post-menopausal cisgender women, we were able to show that androgen therapy induces consistent changes in the breast that supersede those related to the cession of menses and estrogen production by the ovaries. In line with observed reduction of mammary gland tissue in transgender men, we found androgen increased chromatin condensation, general silencing of transcriptional outputs as well as reduced protein translation as inferred through pathway analysis, and some of these changes were also visible in some form when comparing data from cis-male and cis-female breast tissues^11^.

Particularly in luminal-HR^+^ cells most of the identified changes were linked directly to AR regulation. For example the PGR nuclear receptor, which regulates breast development, the menstrual cycle and lactation, was strongly downregulated and showed chromatin-based evidence of AR binding at its gene body^22^. Here we also observed increased activity of FOXA1 and NFIC, two transcription factors that have been shown to be co-regulators for AR in breast and prostate cancer, thus revealing similarities to regulatory mechanism seen in other tissues^22, 92, 93^. In contrast to PGR silencing, we observed AR induced upregulation of *CUX2*, a transcription factor found highly expressed in the prostate, and a general shift of gene expression towards programs seen in cis-male breast. One consequence of this transcriptional reprogramming is altered secretion of ligands, such as AZGP1 and the mammary growth factors AREG, and EREG, enabling luminal-HR^+^ cells to communicate hormonal changes to neighboring cell types lacking AR expression. Related to this, we found that myoepithelial basal cells of the breast lost expression of *OXTR* and *ACTA2*, which are essential for duct contraction during lactation^47, 100^. This also corresponds to reports that breastfeeding rate is lower in cisgender women with PCOS and that androgen and its derivatives can inhibit physical lactation^47, 101–103^. Overall, this study showed that gender affirming testosterone therapy has the intended consequence of inducing cis-male-specific changes in the breast to better align this organ with the gender identify of transgender men.

One aspect of the changes induced that we were not able to fully elucidate mechanistically, was a reduction of estrogen signaling programs in luminal-HR^+^ cells. *ESR1* expression was not altered in transgender men, nor was overall chromatin accessibility of binding motifs for this nuclear receptor. Instead, we inferred from altered combinatorial motif binding that estrogen-signaling related changes were specific to genes that are co-regulated by ESR1 and AP-1 factors, specifically JUN. This could indicate the decreased affinity of ESR1 towards JUN or decreased availability of JUN for co-regulation of genes with ESR1 upon androgen treatment, presumably through redirection by AR. As a result, changes in ESR1-related pathways could, at least partially, occur due to decreased activity of AP-1 factors as well. Likely related to this, one other study has shown that blocking AP-1 factors overcomes endocrine therapy resistance^104^. In future studies, it will be important to incorporate AR, ESR1 and JUN ChIP-seq data and functional studies to determine the mechanisms through which estrogen signaling is silenced by AR in these normal breast cells, and the role that AP-1 transcription factors play.

Since ER^+^ breast cancer proliferation is thought to be driven by estrogen signaling and since the rate of breast cancer is lower in cisgender and transgender men, we also examined cancer-related pathways and how they respond to androgen^8, 105^. Among the three epithelial cell types, luminal cells correlated most strongly with PAM50 classifications of breast cancer, while basal cells did not present with a significant association (Fig. S13A). Luminal-HR^−^ cells that are believed to give rise to basal-like breast tumors were indeed specifically correlated with the corresponding signature and showed a significant reduction in their similarity after androgen treatment (Fig. S13B)^106, 107^. Additionally, upon treatment, correlation of luminal-HR^+^ cells with luminal A- and B-type breast cancers decreased more than any other comparison of cell types with tumor types, corroborating previous findings that androgen treatment suppresses genes driving hormone receptor positive breast cancers (Fig. S13B)^8^. In this study, Hickey et.al., show androgen suppresses proliferation of estrogen-dependent breast cancer cells, and we found that genes used to make this inference were also altered in transgender male luminal-HR^+^ cells; for example, both studies showed downregulation of *PGR* and the oncogene *BCL2* (apoptosis inhibitor), as well as upregulation of the tumor suppressor *SEC14L2* (Fig. S13C)^108–110^. As mentioned above, our data also shows androgen treatment inhibits secretion of the epithelial growth factors AREG and EREG by luminal-HR^+^ cells as well as their receptor EGFR, which are all overexpressed in ER^+^ breast cancers and implicated in breast cancer tumorigenesis and invasiveness^30–34^.

Global chromatin silencing in the breasts of transgender men may also have a protective effect for breast cancers, and as angiogenesis and nutrient supply are a hallmark of cancer, it is tempting to hypothesize that reduced angiogenic activity on capillary cells in testosterone treated breasts could also contribute to this protection. We also observe changes in the local immune microenvironment of the breast epithelium with macrophages being reduced and CD4 T-Cells showing a specific increase and a migratory response toward epithelial structures. This clearly illustrates the immune modulatory capacity of androgen and suggests that our current understanding of tumor infiltrating lymphocytes (TILs) might benefit from further research on androgen in this field. Furthermore, these findings provide some insight into the biological basis of sexual dimorphism seen in immune responses and susceptibility to autoimmune diseases^111^.

Though we did not perform metabolomic profiling, we found multiple lines of evidence indicating metabolic activity of the breast is affected by androgen, in line with a litany of studies that have shown metabolic changes are induced by sex hormones^77, 112, 113^. The most striking related changes were seen in adipocytes. In line with increased insulin sensitivity observed in transgender men, we saw upregulation of the insulin receptor and correspondingly altered downstream effectors of the PI3K pathway^112^. NR4A1, an orphan nuclear receptor and key regulator of glucose and lipid metabolism was found strongly upregulated in adipocytes and other cell types as well^87, 114^. These findings support ongoing research on NR4A1, indicating it aids in insulin signaling and that it is an important target for the study of diabetes and metabolic disorders, particularly in transgender patients^84–86, 114^. Other mechanisms through which androgen was inferred to affect adipocyte function were through the above-mentioned upregulation of *AZGP1*, and activation of the anti-adipogenic transcription factor *TCF7L2*, with this potentially being an undiscovered mechanism through which transgender men shed weight from other metabolically active fat stores^93, 96, 97^. Future work should include precise metabolic profiling of blood and local tissues in testosterone treated individuals to better understand these metabolic changes and if findings from here could be used to perhaps treat obesity.

The only prior study to perform a broad molecular assessment of breasts from transgender men used gene expression microarray technology to analyze RNA from bulk pre- and post-surgical biopsies of the breast tissue^115^. Despite the difference in fidelity, some of our key results, including the global downregulation of genes, decreased translation, and increased activity of NR4A1 remain consistent in both studies. The single-cell resolution and the multi-modal aspect of our approach, however, allowed us to understand the dynamics of the breast cell type gene regulatory networks at an unprecedented level. Loss of expression and chromatin activity for transcription factors including AP-1 factors, for example, was not detectable in the previous study, possibly due to the heterogeneity of the breast tissue and the cell-type–specific pattern we observed in epithelial cells. Our in-depth spatial profiling efforts also allowed us to better understand how the morphology and local microenvironment of the breast are altered by androgen and allowed further inference of the linkage between gene expression programs in neighboring cell types. Important caveats of this study include low patient numbers, which resulted from a combination of sample availability as well as the cost and complexity of the analyses performed. These numbers and a lack of some relevant information about some samples, such as menstrual status, systemic hormone levels and body fat composition prevented us from elucidating how these major contributors to breast tissue homeostasis impact the response to androgen. Overall, we have developed a rich resource for the research community to study hormonal control of human tissue homeostasis, while also providing a roadmap for future studies aiming to study molecular and cellular changes brought on by gender-affirming therapies.

## Supporting information

Table.1-CellType_Markers

Table.2-Cohort_Details

Table.3-Subcluster_Marker_Genes

Table.4-CellType_DE_Response

Table.5-Pathway_Enrichments

Table.6-gProfiler_Results

Table.7-Antibodies_CODEX_tissue_microarray

## Supplementary Figures

**Supplementary Figure S1:**
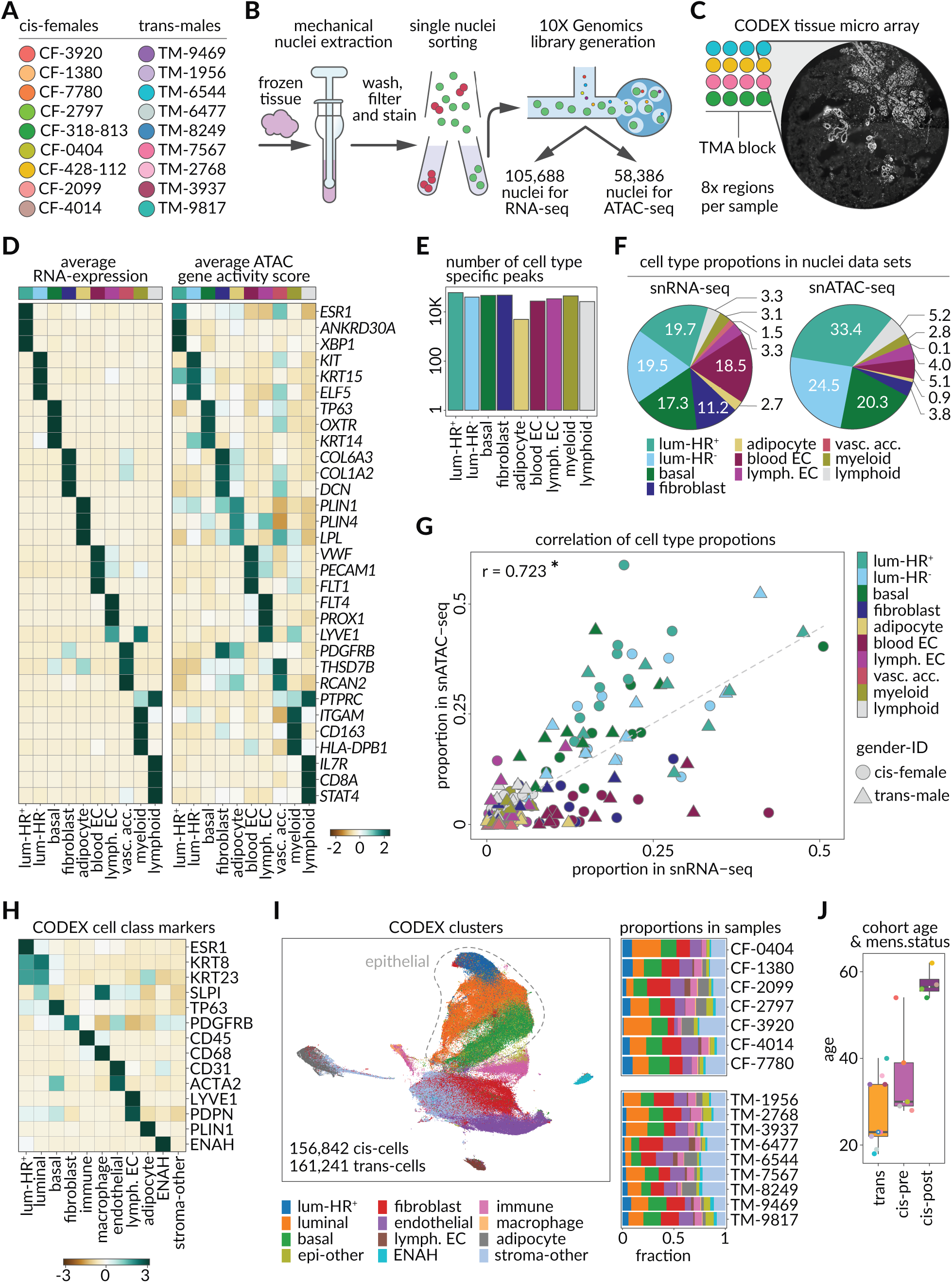
Sample preparation and experimental design. **A)** Anonymized patient IDs and their color code used for data display. **B)** Schematic of key processing steps of snRNA-seq and snATAC-seq library generation. Nuclei were released from cryopreserved mammary tissue through mechanical homogenization, sorted to enrich for singlets and directly used as input for 10X Genomics library preparation. **C)** Schematic design of 4×4 tissue microarrays used in CODEX experiments, colors indicating sample arrangement. We used 7 cis-female and 9 trans-male samples, using 8 (2 mm) tissue regions from left and right breast of each sample. We block-randomized the samples so that each TMA contains both cis-female and trans-male samples. **D)** Heatmap shows the markers we used to identify cell types in snRNA and snATAC data. Values indicate scaled averages of RNA-expression in snRNA-seq (left) and gene activity scores in snATAC-seq data sets (right). **E)** Number of cell type specific chromatin accessibility peaks within each cell type. **F)** Relative proportion of each cell type within snRNA-seq (left) and snATAC-seq data (right). **G)** Scatterplot shows the fraction of each cell type within each of the 18 samples in snRNA-seq data (horizontal axis) and snATAC-seq data (vertical axis). The dashed diagonal represents the identity line (p-value = 1.85 x 10^-28^). **H)** Heatmap shows the scaled average staining intensities corresponding to each of the cell class markers within each group of cells in the CODEX tissue microarray dataset. **I)** UMAP shows identity of cells in CODEX tissue microarray dataset determined through clustering based on staining intensities of cell class markers. Barplots show the proportion of each cell class within each of the 16 samples we used for CODEX tissue microarray. **J)** Boxplot shows age of the trans-male samples (orange), cis-female pre-menopausal samples (light purple), and cis-female post-menopausal samples (dark purple).

**Supplementary Figure S2:**
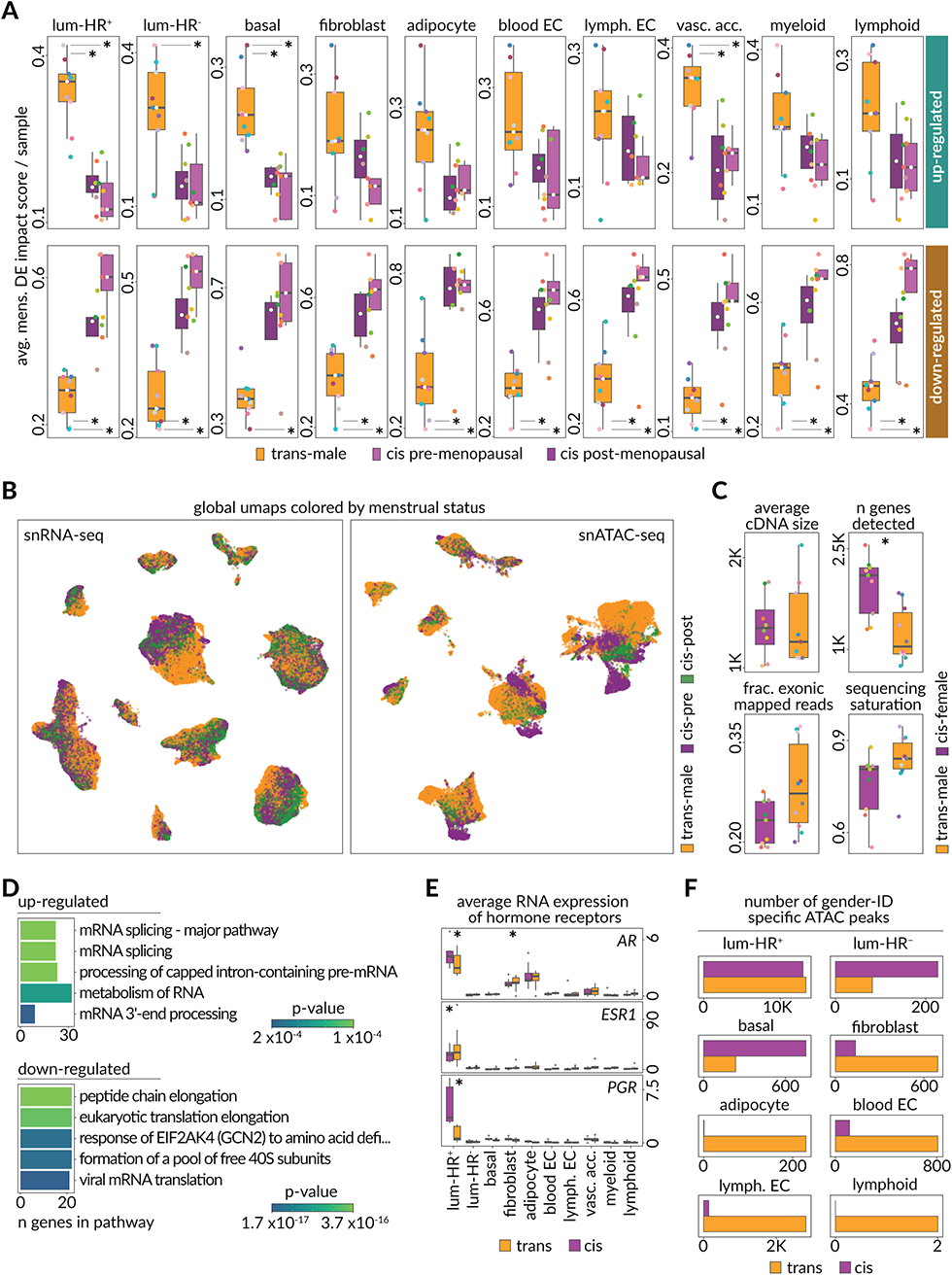
Global transcriptional differences of cell types. **A)** Boxplots showing average differential expression impact scores for each sample organized by cell types and menstrual status (trans-male = orange, pre-menopausal cis-female = light purple, post-menopausal cis-female = dark purple). Differentially expressed genes with an avg. log_2_FC > 0.5 (upregulated) or < −0.5 (downregulated) between trans-male and cis-female in each cell type were used to score all cells, providing a metric on how much a sample contributes to the observed differential expression. Significant differences are marked by (*). **B)** UMAPs show individual nuclei extracted from trans-male (orange), pre-menopausal cis-female (purple), and post-menopausal cis-female (green) breast tissues in snRNA-seq (left) and snATAC-seq (right) datasets. **C)** Boxplots show average size of cDNA in base pairs as measured by BioAnalyzer during library preparation (top left; p-value, Wilcoxon = 0.89), median number of genes as reported by CellRanger (top right; p-value, Wilcoxon = 0.0041), fraction of reads mapped to exonic regions as reported by CellRanger (bottom left, p-value, Wilcoxon = 0.066), and sequencing saturation as reported by CellRanger (bottom right; p-value, Wilcoxon = 0.065) in cis-female (purple) and trans-male (orange) samples. **D)** Pathway enrichment of globally up- and downregulated genes. Genes found differentially expressed in at least 9 of 10 cell types were enriched for Reactome pathways. Five most significant results are shown for up- and downregulated genes in trans-male breast tissues. **E)** Sample averages of RNA-expression levels of sex hormone receptors for androgen (*AR*), estrogen (*ESR1*) and progesterone receptor (*PGR*) across all identified cell types. (adjusted p-values: luminal-HR+, MAST: AR = 4.17 x 10^-170^, ESR1 = 2.95 x 10^-172^, PGR < 2.95 x 10^-172^, adipocyte: *AR* = 5.67 x 10^-12^).Barplots show the number of chromatin accessibility peaks specific to trans-male cells (orange) or cis-female cells (purple) within each snATAC-seq cell type.

**Supplementary Figure S3:**
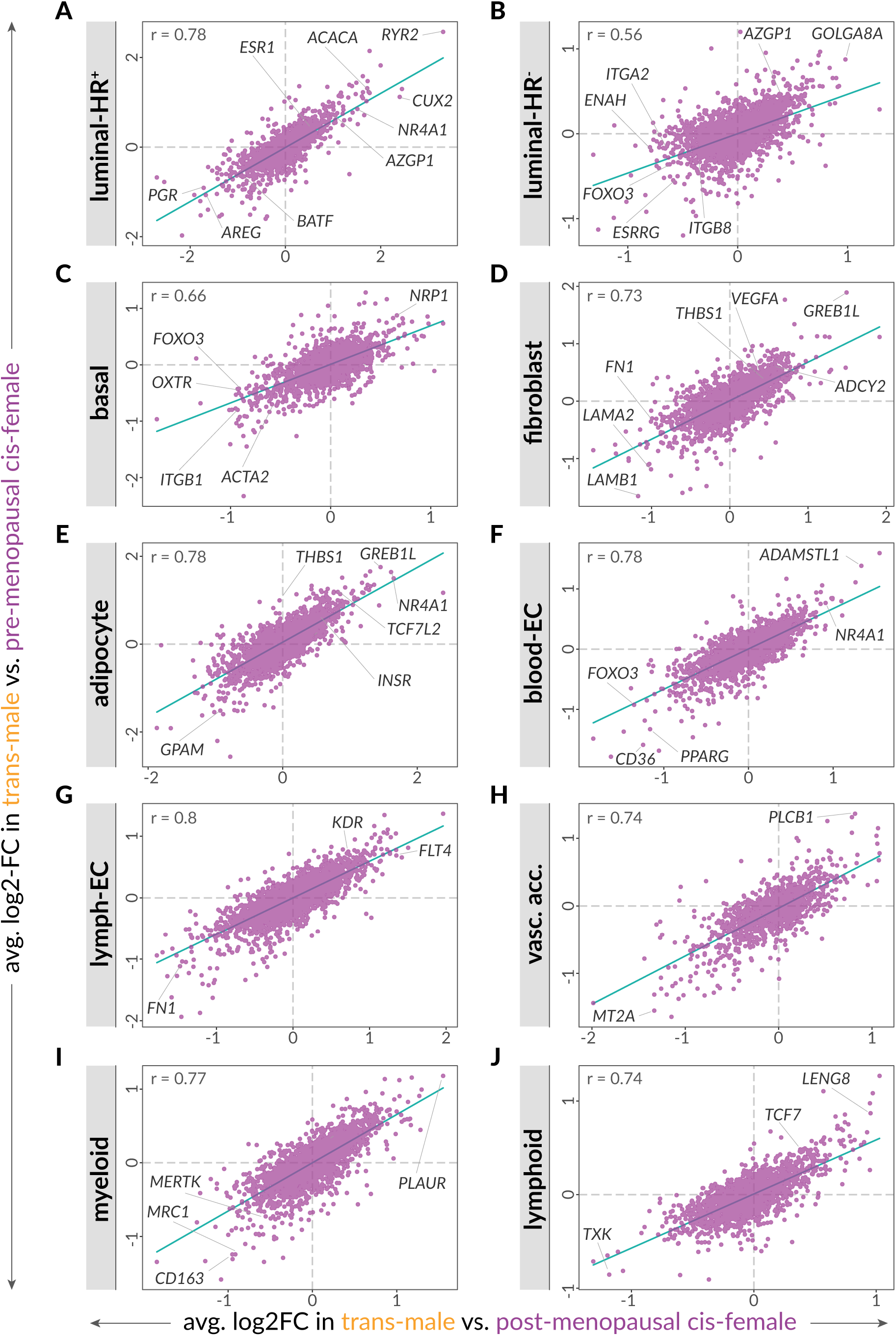
Similarity of pre-menopausal and post-menopausal cis-female cells when comparing their transcriptome to transgender male cells. **A)** Horizontal axis shows the average log_2_ fold change in expression when comparing trans-male luminal-HR^+^ cells to post-menopausal cis-female luminal-HR^+^ cells. The vertical axis shows the average log_2_ fold change in expression when comparing trans-male luminal-HR^+^ cells to pre-menopausal cis-female luminal-HR^+^ cells. Annotations represent some key differentially expressed genes in luminal-HR^+^ cells discussed in the study. **B–J)** Similar to (**A**) for the other 9 cell types (p-values for all 10 plots < 2.22 x 10^-16^).

**Supplementary Figure S4:**
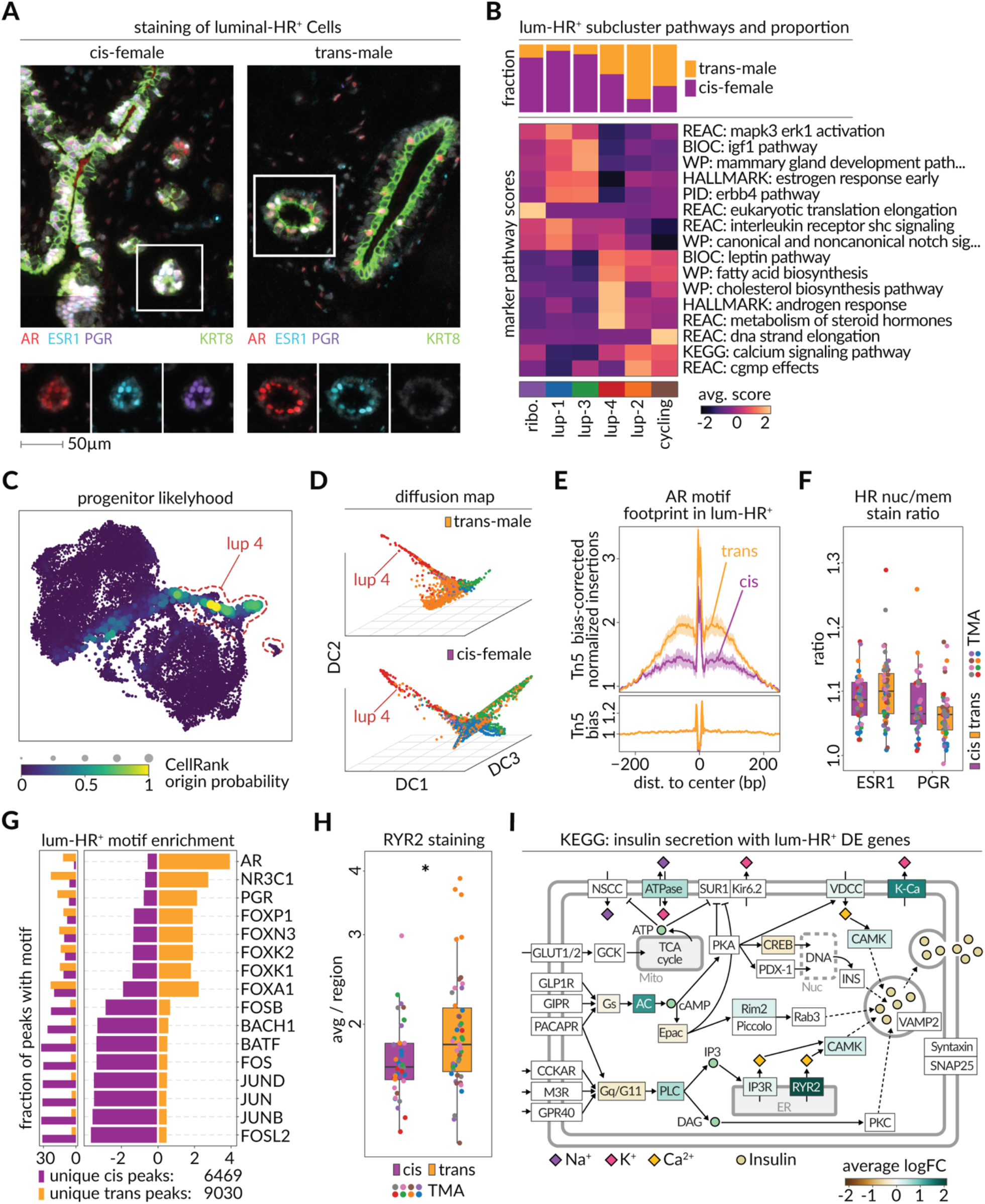
Regulatory dynamics of transgender hormone-responsive epithelial cells. **A)** Representative CODEX microarray images, showing hormone receptor staining (AR = red, ESR1 = blue, PGR = purple) among luminal epithelial (KRT8 = green) cells in cis-female (left) and trans-male (right) breast tissues. Both tissue regions are part of the same TMA. **B)** Heatmap shows average module scores of biological pathways with differential enrichment among the 6 subclusters of luminal-HR^+^ cells. The top bar plot indicates the fraction of each subcluster among either trans-male (orange) or cis-female (purple) cells. **C)** Origin probability, as calculated by CellRank, among luminal-HR^+^ cells overlaid in the corresponding UMAP from Fig. 2A. Location of cells belonging to subcluster lup 4 is highlighted in red. **D)** Diffusion map of trans-male (top) and cis-female (bottom) luminal-HR^+^ cells, generated with destiny. Colors represent subclusters from Fig. 2A and subcluster lup 4 is labeled. **E)** Motif footprints for Androgen Receptor (AR, AR-CisBP M03389_2.00) among trans-male (orange) and cis-female (purple) luminal-HR^+^ cells. Top panel shows the transposase bias-corrected signal and the bottom panel shows the transposase bias. **F)** Ratio of nuclear to cytoplasmic staining signal of hormone receptors ESR1 and PGR in cis-female (purple) and trans-male (orange) tissues on CODEX microarrays. Dot colors indicate from which TMA the shown regions originate. **G)** Right panel shows the enrichment of motifs among the peaks of trans-male (orange) and cis-female (purple) luminal-HR^+^ cells. Left panel shows the fraction of the peaks of the corresponding cells which overlap with the motif. **H)** Average RYR2 staining intensity per tissue region of luminal-HR^+^ cells in cis-female (purple) and trans-male (orange) tissue of CODEX microarray data. Dot colors indicate the origin TMA of shown regions (p-value, Wilcoxon = 0.024). **I)** KEGG Insulin secretion pathway annotated with average log_2_ fold change expression of the involved genes. Higher values indicate a higher expression in trans-male luminal-HR^+^ cells.

**Supplementary Figure S5:**
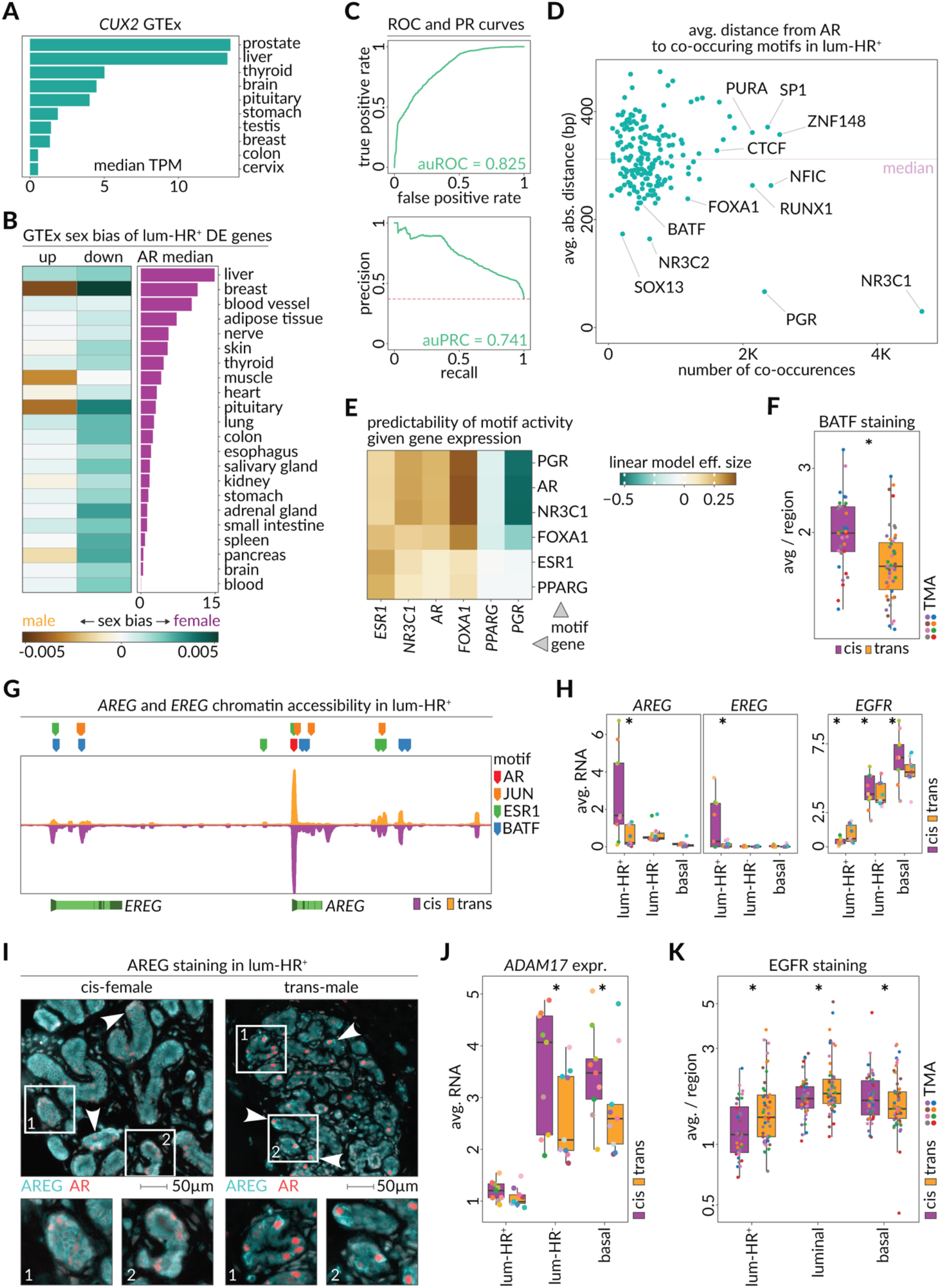
Downstream effects of regulatory changes in transgender luminal-HR^+^ cells. **A)** Gene expression of *CUX2* transcription factor among the top 10 *CUX2*-expressing tissues of the GTEx dataset. The horizontal axis shows median transcripts per million reads (TPM) among the different samples of each tissue. **B)** The heatmap shows the average sex-bias (as provided by GTEx) of upregulated (trans-male–specific) and downregulated (cis-female–specific) genes (|log_2_FC| = 0.5) of luminal-HR^+^ cells among the cis-male or cis-female samples of each tissue. The figure shows the tissues with data in both cis-male and cis-female samples. Ordering of tissues represents their median *AR* gene expression. **C)** Receiver-operating characteristic curve (top) and precision-recall curve (bottom) assess the performance of the random forest on the held-out set of peaks. **D)** Average motif distance and total number of co-occurrences between AR and other transcription factor motifs in open chromatin regions of luminal-HR^+^ cells. Purple horizontal dashed line indicates the median average motif distance. **E)** The heatmap shows the effect size of linear models which predict a transcription factor motif (vertical axis) z-score activity using the gene expression of the transcription factor (horizontal axis). **F)** Boxplots show per-region average BATF staining intensity in CODEX microarray data of cis-female (purple) and trans-male (orange) tissues. Dot colors indicate TMA origin of the shown regions. (p-value, Wilcoxon = 0.00014) **G)** Selected transcription factor binding sites (colored markers) in open chromatin regions around *EREG* and *AREG* in luminal-HR^+^ cells. Transcription factors were selected based on differential expression and motif enrichment. The genomic window shows chromatin accessibility for trans-male (orange) and cis-female (purple) cells. Gene bodies of *EREG* and *AREG* (light-green) are shown with exon boundaries (dark green) and promoter location (arrow). **H)** Average RNA expression of *AREG*, *EREG*, and their receptor *EGFR* among luminal-HR^+^, luminal-HR^−^, and basal epithelial cells of cis-female (purple) and trans-male (orange) samples (adjusted p-values, MAST: *AREG* in luminal-HR^+^ < 2.2 x 10^-16^, *EREG* in luminal-HR^+^ < 2.2 x 10^-16^, *EGFR* in luminal-HR^+^ = 4.37 x 10^-107^, luminal-HR^-^ = 2.94 x 10^-205^, basal = 8.18 x 10^-117^) **I)** Microscopic images from CODEX microarray data showing AREG staining (turquoise) and luminal-HR^+^ cells (AR = red) in cis-female (left) and trans-male (right) tissues, each with arrows and zoomed-in panels to highlight AREG distribution around luminal-HR^+^ cells. Both tissue regions are part of the same TMA. **J)** Left panel boxplots show per-sample average expression of *ADAM17* among basal, luminal-HR^−^, and luminal-HR^+^ cells of cis-female (purple) and trans-male (orange) (adjusted p-values, MAST: luminal-HR^-^ < 2.2 x 10^-16^, basal = 1.32 x 10^-265^). **K)** Boxplots show per-region average EGFR staining intensity of epithelial cell types in CODEX microarray data of cis-female (purple) and trans-male (orange) tissues. Dot colors indicate TMA origin of the shown regions. (p-value, Wilcoxon: luminal-HR^+^ = 0.0069, luminal = 0.083, basal = 0.26)

**Supplementary Figure S6:**
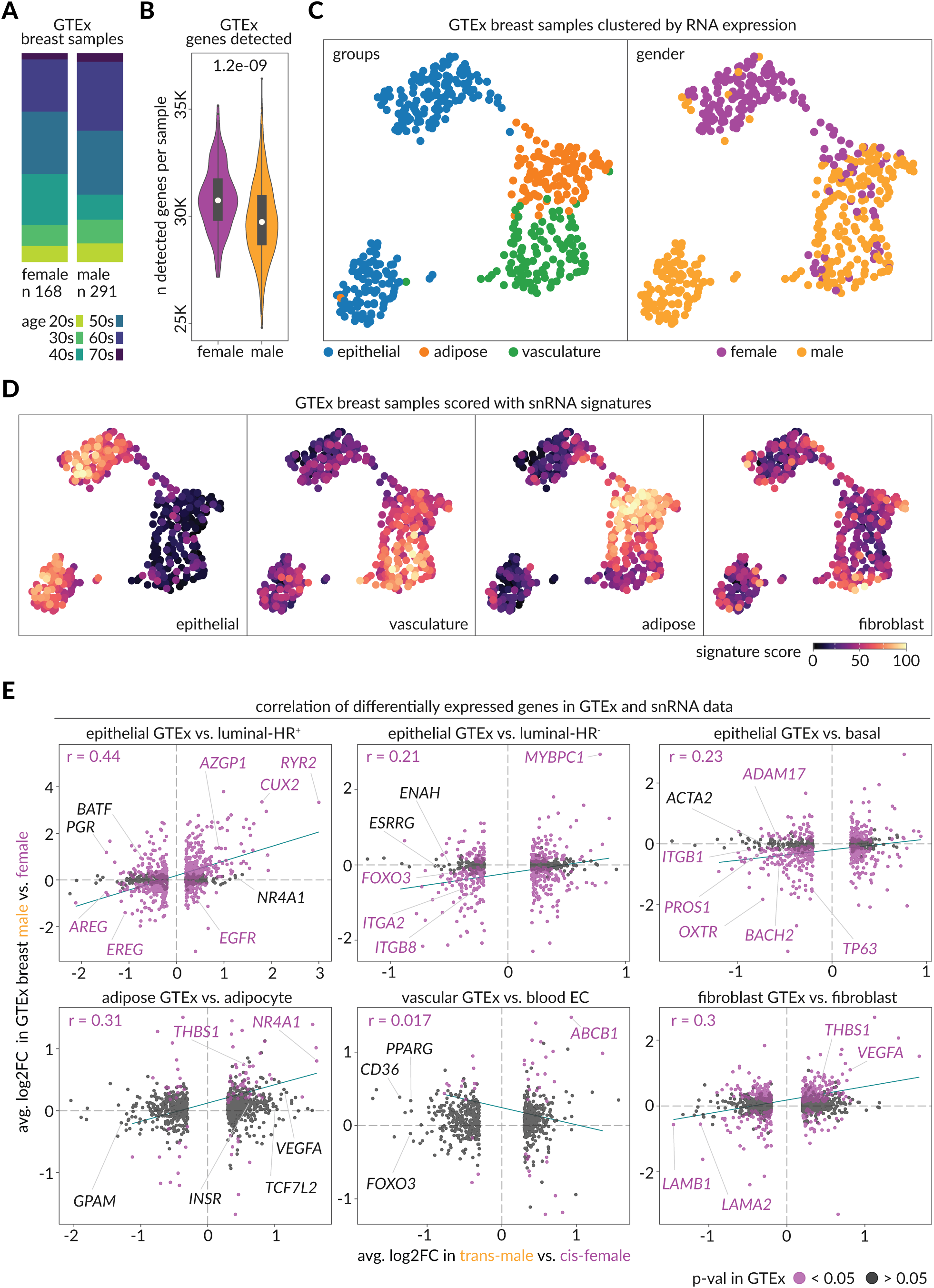
Transcriptome similarity of cis-male, cis-female, and trans-male breast tissues. **A)** Barplot shows age groups (color) of and number of cisgender men (left) and cisgender women (right) GTEx donors of breast tissue. **B)** Number of genes detected in GTEx breast samples among cis-female (purple) and cis-male (orange) samples. **C)** Left panel shows UMAP of GTEx breast tissues colored according to their similarity to snRNA gene signatures of epithelial (blue), adipose (orange), or green (vasculature). Right panel shows the same data according to the sex of the tissue donor. **D)** Signatures obtained from snRNA cell types for epithelial, vasculature, adipose, and fibroblast cell types overlaid on the UMAP of GTEx breast samples. **E)** Horizontal axis shows log_2_ fold change of significantly differentially expressed genes comparing trans-male to cis-female cells in cell types of snRNA-seq data. Vertical axis shows log_2_ fold change of genes comparing cis-male to cis-female GTEx breast samples. Purple data points indicate significance in GTEx comparison. Panels correspond to luminal-HR^+^, luminal-HR^−^, basal, adipose, blood endothelial, and fibroblast cells.

**Supplementary Figure S7:**
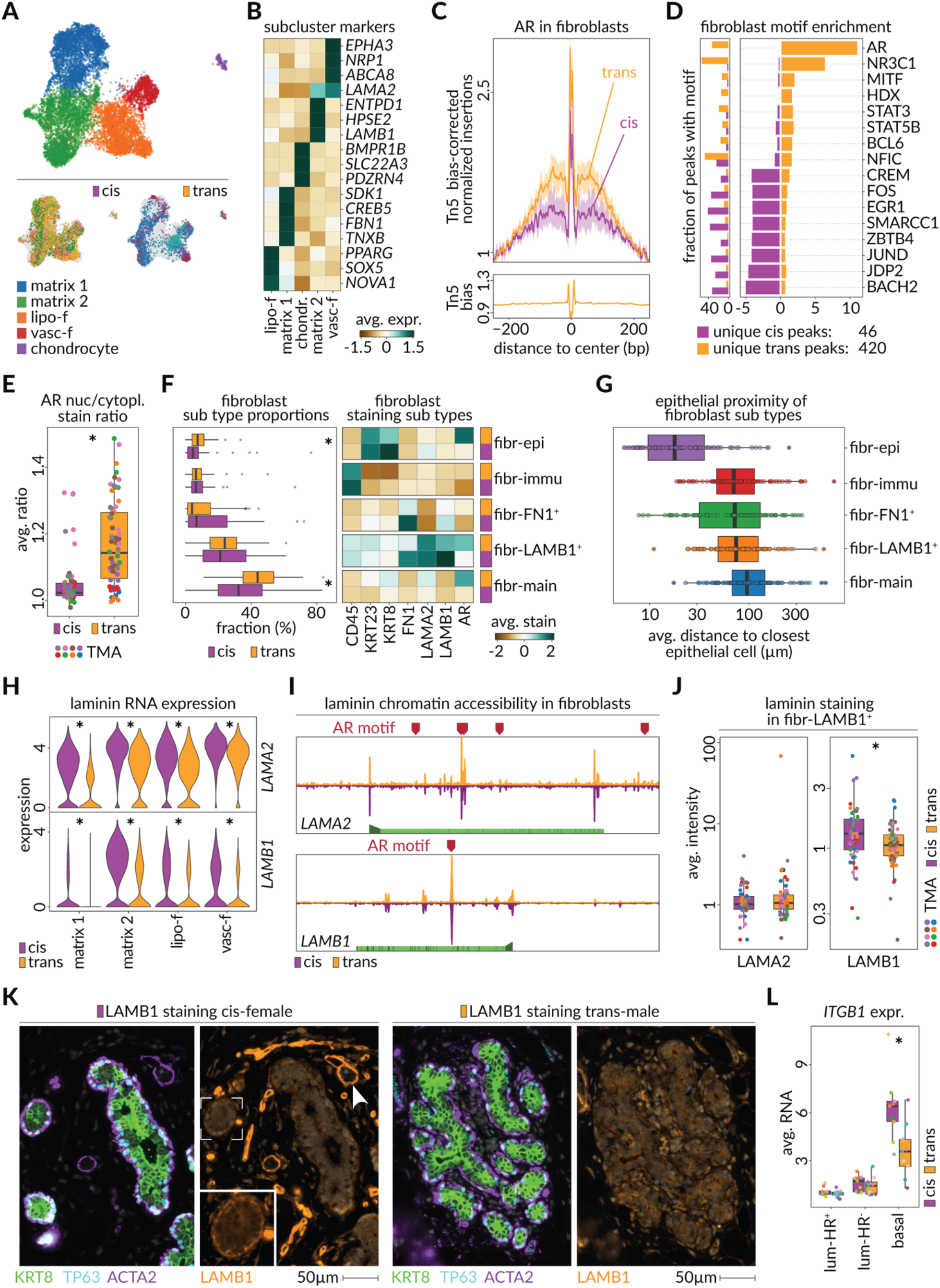
Androgen receptor activity drives fibroblast’s response to hormone replacement therapy. **A)** UMAP depicting fibroblast subclusters (top) in snRNA-seq data (matrix 1 and 2 = matrix-type fibroblasts, lipo-f = lipo-fibroblasts, vasc-f = vascular-like fibroblasts) and the distribution of patient samples in each gender identity (bottom). **B)** Heatmap shows scaled average expression of snRNA-seq markers identified for each of the fibroblast subclusters. **C)** Motif footprints for Androgen Receptor (AR, AR-CisBP M03389_2.00) among trans-male (orange) and cis-female (purple) fibroblast cells. Top panel shows the transposase bias-corrected signal, and the bottom panel shows the transposase bias. **D)** Right panel shows the enrichment of motifs among unique accessible chromatin peaks of trans-male (orange) and cis-female (purple) fibroblast cells. Left panel shows the fraction of the peaks of the corresponding cells which overlap with the motif. **E)** Boxplot shows the average ratio of androgen receptor (AR) staining intensity in fibroblast nucleus compared to the cytoplasm in tissue regions of cis-female (purple) and trans-male (orange) samples in CODEX microarray data (p-value, Wilcoxon = 1.2 x 10^-8^). Dot colors indicate TMA origin of the shown regions **F)** Left panel boxplots show the fraction of cis-female (purple) and trans-male (orange) cells corresponding to 5 different classes of fibroblasts detected in tissue regions of CODEX microarray data (p-values, Wilcoxon: fibr-main = 0.00011, fibr-epi = 0.0046). Right panel shows the scaled staining intensities of various markers that distinguish the 5 subtypes of fibroblasts. **G)** Boxplots show per region average distance for each of the 5 subtypes of fibroblasts to the most proximal epithelial cell. **H)** Violin plots show the RNA expression of laminins *LAMA2* (top) and *LAMB1* (bottom) in fibroblast subclusters, split by cis-female (purple) and trans-male (orange) cells (adjusted p-values, MAST: *LAMA2* in lipo-f = 2.60 x 10^-63^, matrix 1 = 5.11 x 10^-175^, matrix 2 = 1.03 x 10^-69^, vasc-f = 1.99 x 10^-10^; *LAMB1* in lipo-f = 3.71 x 10^-35^, matrix 1 = 1.46 x 10^-55^, matrix 2 = 3.08 x 10^-87^, vasc-f = 2.58 x 10^-13^). **I)** AR binding sites (red markers) across genomic regions of *LAMA2* and *LAMB1*. Gene bodies are shown (light-green) with the promoter (arrow) and exon boundaries (dark-green). Genomic window shows chromatin accessibility in cis-female (purple) and trans-male (orange) fibroblasts. **J)** Boxplots show per-region average LAMA2 (left) and LAMB1 (right) staining intensities in the LAMB1^+^ fibroblast subtype on CODEX microarray data of cis-female (purple) and trans-male (orange) tissues. Dot colors indicate TMA origin of the shown regions. (p-value, Wilcoxon: LAMA2 = 0.64, LAMB1 = 0.0079) **K)** Microscopic images from CODEX microarray data show staining of LAMB1 (orange) among cis-female (left) and trans-male (right) tissues. Other markers include KRT8 (pan-luminal), TP63 (basal cells), and ACTA2 (basal cells). Arrow indicates distinctive LAMB1 ECM structures near epithelium and box highlights LAMB1 layer around acinus. **L)** Boxplots show the average RNA expression of *ITGB1* among luminal-HR^+^, luminal-HR^−^, and basal epithelial cells of cis-female (purple) and trans-male (orange) samples (adjusted p-value, MAST in basal < 2.22 x 10^-16^).

**Supplementary Figure S8:**
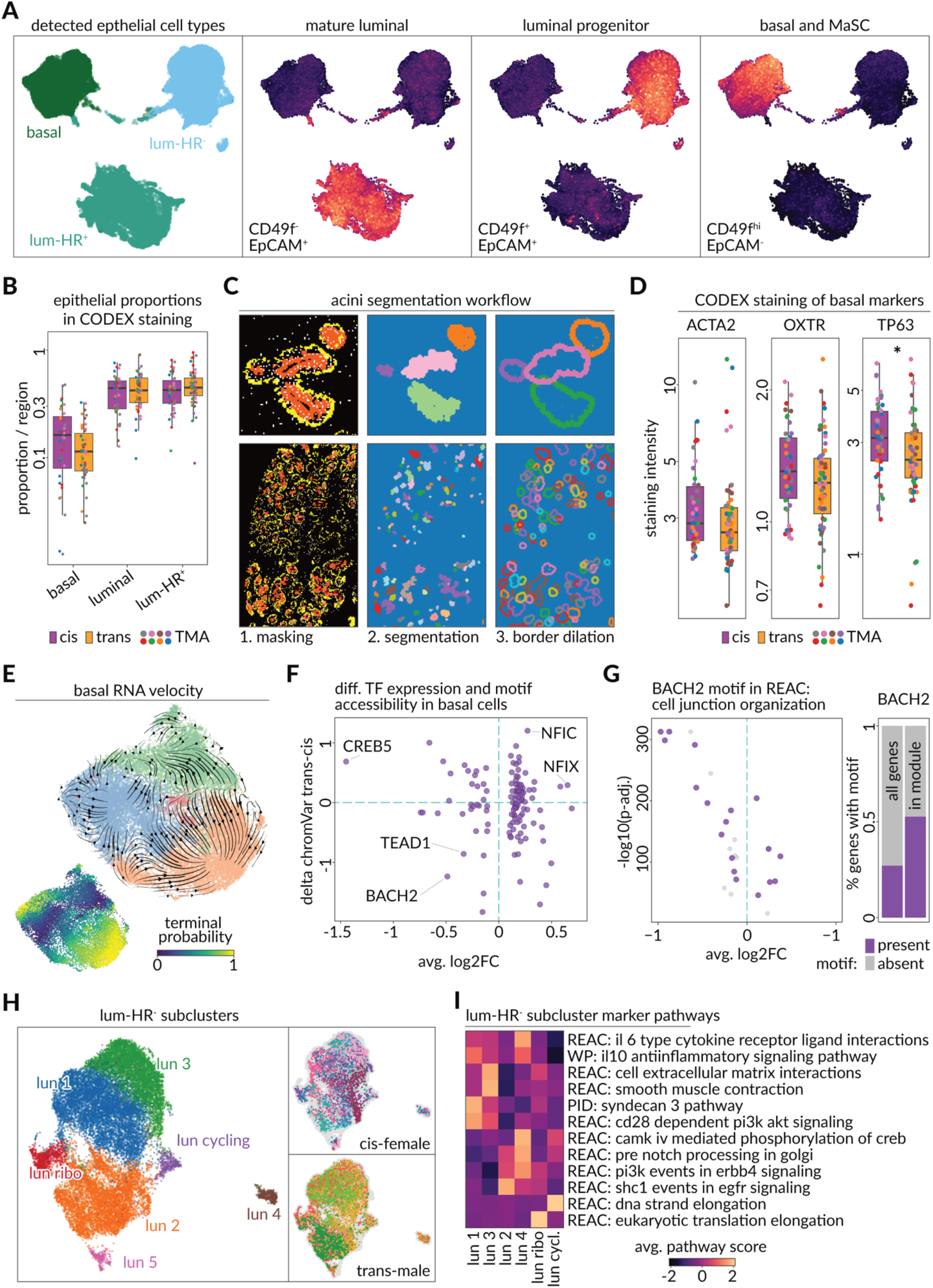
Luminal breast duct reorganization due to changes in basal cell contacts. **A)** UMAPs show RNA-seq modules from (Lim et.al., 2009) overlaid on epithelial cells of snRNA-seq data. The corresponding flow cytometry markers used for detecting them are shown on the bottom. **B)** Boxplots show the fraction of each of the three epithelial cell types (basal, luminal, and luminal-HR^+^) within each region of CODEX microarray data in cis-female (purple) and trans-male (orange) tissues. Dot colors indicate TMA origin of shown regions. **C)** The process of segmenting acini structures through masking luminal and basal cells (orange = KRT8 + KRT23, yellow = ACTA2 in left panel, respectively), segmenting the image, and expanding boundaries to estimate coverage of the smooth muscle layer. **D)** Boxplots show the average staining intensity of ACAT2, OXTR, and TP63 proteins among basal cells in tissue regions of cis-female (purple) and trans-male (orange) samples. (p-values, Wilcoxon: ACTA2 = 0.054, OXTR = 0.16, TP63 = 0.04) **E)** UMAP shows RNA-velocity streams over subclusters of basal cells (top) and the overlaid CellRank probability of terminal state (bottom). **F)** Scatterplot shows the log_2_ fold change in expression of a given transcription factor (horizontal axis) and the difference in chromVAR z-score of the transcription factor (vertical axis) when comparison trans-male to cis-female basal cells. **G)** Volcano plot shows basal cell log_2_ fold change in expression (horizontal axis) and −log_10_ p-value (vertical axis) of genes annotated within the Reactome database pathway for cell junction organization. Purple data points contain a chromatin accessibility peak with a BACH2 transcription factor sequence motif (CisBP BACH2_113) match. Barplot shows the fraction of all genes (left) and Reactome pathway for cell junction organization genes (right) which contain a chromatin accessibility peak containing a BACH2 sequence motif match. **H)** UMAP plots of luminal-HR^−^ cells in RNA space, showing detected subclusters (left) and distribution across cis-female and trans-male samples (right). **I)** Heatmap shows enrichment of luminal-HR^−^ subclusters in marker pathways. Values show scaled average module scores. For each subcluster, the two most significant differentially expressed pathways were selected. (REAC = Reactome, WP = WikiPathways, BIOC = Biocarta, KEGG = Kyoto Encyclopedia of Genes and Genomes, PID = Pathway Interaction Database).

**Supplementary Figure S9:**
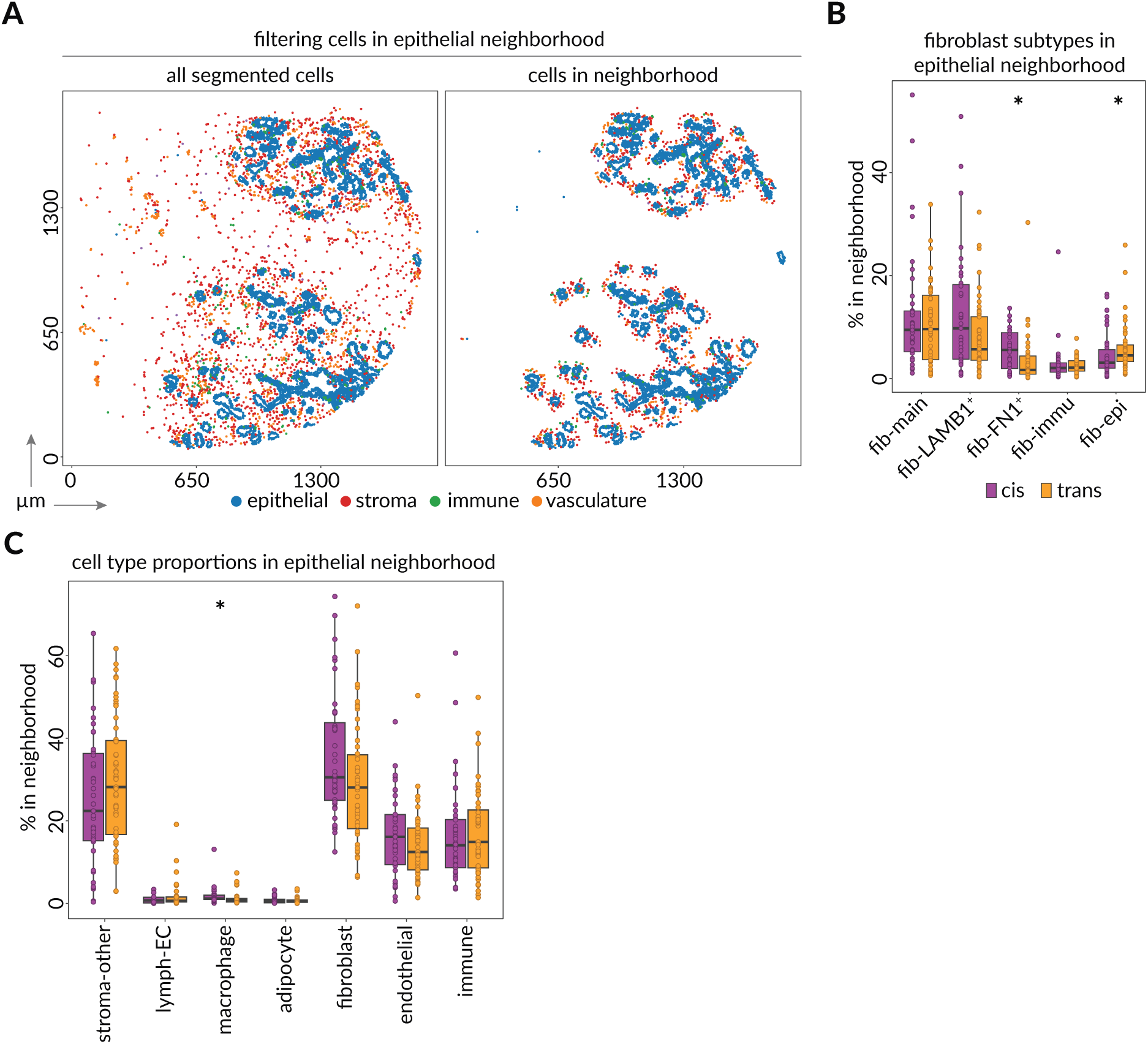
Stromal composition of epithelial neighborhood is altered after androgen treatment. **A)** Schematic shows annotated epithelial (blue), stromal (red), immune (green), and vasculature (orange) cells annotated within a microscopic image from the tissue microarray data. Right shows filtered cells used in neighborhood proximity analyses of epithelial cell types. **B)** Boxplot shows proportions of the 5 fibroblast subtypes among the cis-female (purple) and trans-male (orange) epithelial neighborhoods. **C)** Boxplots show the proportion of each cell type within the epithelial neighborhood of cis-female (purple) and trans-male (orange) samples.

**Supplementary Figure S10:**
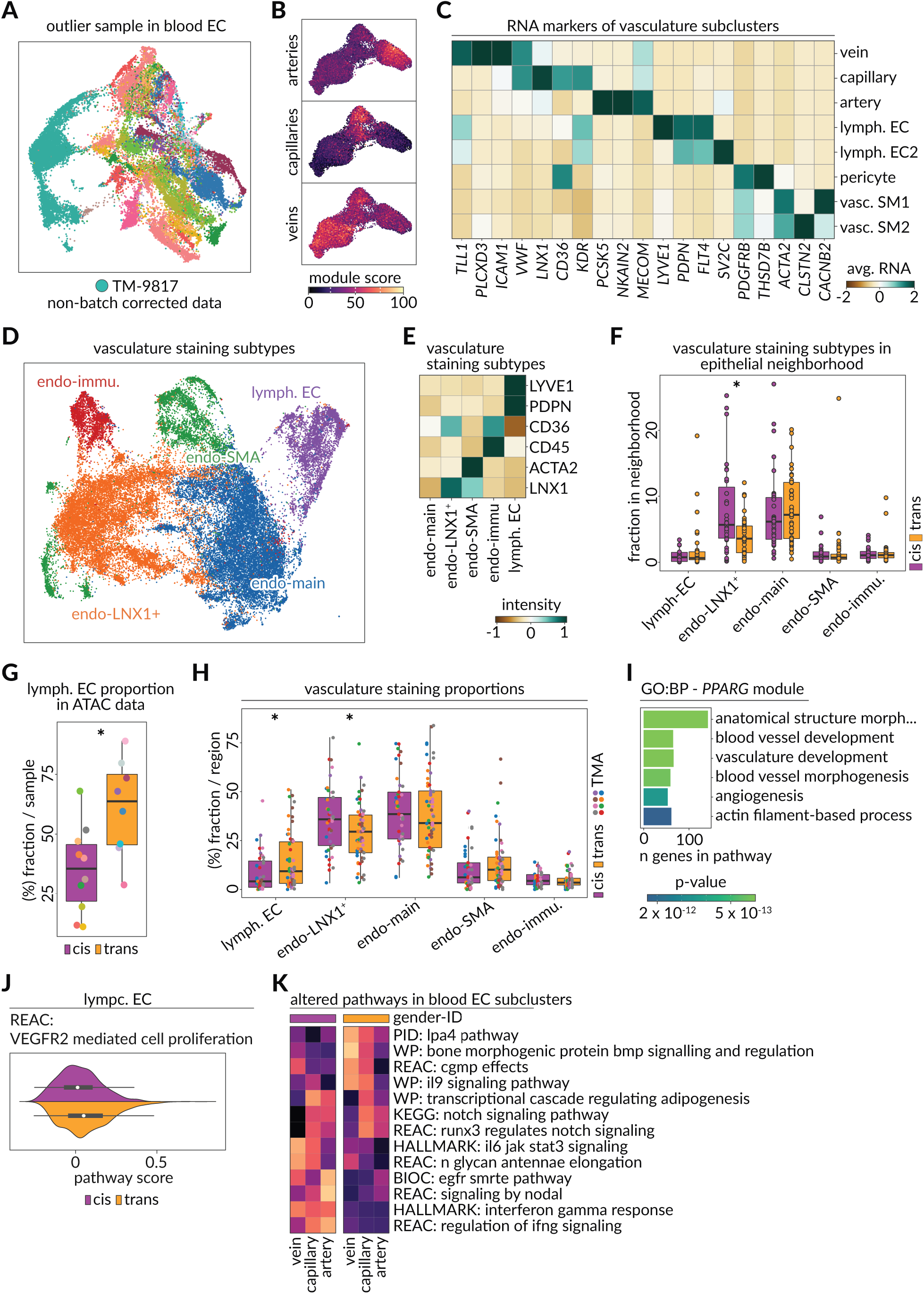
Transcriptome and spatial dynamics of breast vasculature upon hormone replacement therapy. **A)** UMAP shows the sample identity of blood endothelial cells among the 18 samples without applied batch correction. Sample TM-9817 was found to be a strong outlier and was excluded from vasculature analysis. **B)** Gene modules from (Kalucka et.al., 2020) were used to define arteries (top), capillaries (middle), and veins (bottom) in snRNA-seq data. Scores are overlaid on the UMAP plot of blood endothelial cells. **C)** Heatmap shows column Z-score for average expression of the vasculature subcluster markers among the different subtypes of vasculature. **D)** UMAP shows staining subtypes of vasculature cells as determined from CODEX tissue microarray data. **E)** Heatmap shows row Z-score of vasculature marker staining score among the 5 staining subtypes of vasculature cells in CODEX tissue microarray data. **F)** Boxplot shows the fraction of different vasculature subtypes within the neighborhood of epithelial cells in cis-female (purple) and trans-male tissues. **G)** Boxplots show the fraction of lymphatic endothelial cells within snATAC-data of cis-female (purple) and trans-male (orange) samples (p-value, Wilcoxon = 0.021). **H)** Boxplots show the fraction of the different staining subtypes of vasculature in cis-female (purple) and trans-male (orange) tissue regions of CODEX microarray data. (p-values, Wilcoxon: lymph. EC = 0.019, endo-LNX1^+^ = 0.037, endo-main = n.s., endo-SMA = n.s., endo-immu. = n.s.) **I)** Barplot shows the most significantly enriched biological processes (BP) of the *PPARG* module in blood ECs. GO enrichment was done on the 95th percentile of *PPARG* co-expressed genes using g:Profiler. **J)** Kernel density estimate and boxplot of the Reactome VEGFR2-mediated cell proliferation pathway score in cis-female (purple) and trans-male (orange) lymphatic endothelial cells. (p-value, Wilcoxon = 1.06 x 10^-16^) **K)** Selected significantly altered pathways among subclusters of blood endothelial cells are shown as pathway module scores of trans-male and cis-female cells.

**Supplementary Figure S11:**
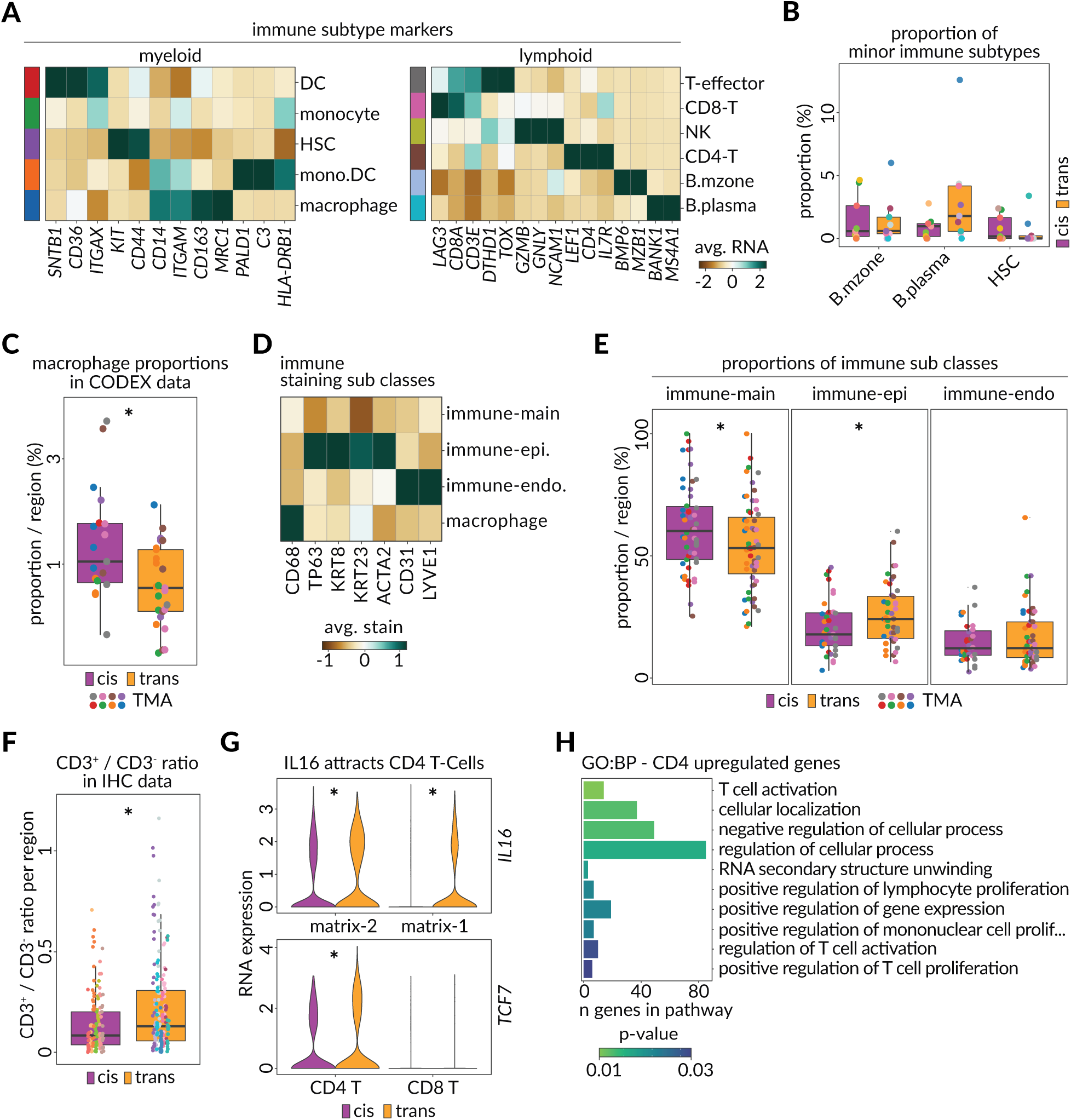
Transcriptome and spatial transition of the breast immune cells upon hormone replacement therapy. **A)** Heatmaps show scaled average RNA expression for markers of myeloid (left) and lymphoid (right) subclusters in the immune compartment. **B)** Boxplots show the proportion of the minor immune subclusters (B-cells and hematopoietic stem cells) among the immune cells of cis-female (purple) and trans-male (orange) samples. **C)** Boxplot shows the proportion of macrophages detected in CODEX microarray regions of cis-female (purple) and trans-male (orange) tissues. Only regions with at least 15 detected macrophages are included. Dot colors indicate which TMA the regions belong to. (p-value, Wilcoxon = 0.011) **D)** Heatmap showing scaled average staining intensity of immune cell markers among the 4 spatial subclasses of immune cells in CODEX microarray data. **E)** Boxplots show the proportion of immune cell spatial subclasses in cis-female (purple) and trans-male (orange) tissue regions of CODEX microarray data. (p-values, Wilcoxon: immune-main = 0.043, immune-epi. = 0.048, immune-endo = n.s.) **F)** Boxplot shows the ratio of immune cells (CD45^+^) expressing CD3 to those not expressing CD3 in IHC scan-regions in cis-female (purple) and trans-male (orange) tissues (p-value, Wilcoxon = 0.0014). **G)** Violin plots show the RNA expression of *IL16* in matrix-2 and matrix-1 fibroblasts (top) and the RNA expression of *TCF7* in CD4^+^ and CD8^+^ T lymphocytes among cis-female (purple) and trans-male (orange) cells. (adjusted p-values, MAST: *IL16* in matrix-1 = 3.81 x 10^-10^, matrix-2 = 2.66 x 10^-01^, *TCF7* in CD4 = 6.85 x 10^-08^, CD8 = n.s.) **H)** Barplot shows the most significantly enriched biological processes (BP) of genes upregulated in transgender male CD4+ T-cells.

**Supplementary Figure S12:**
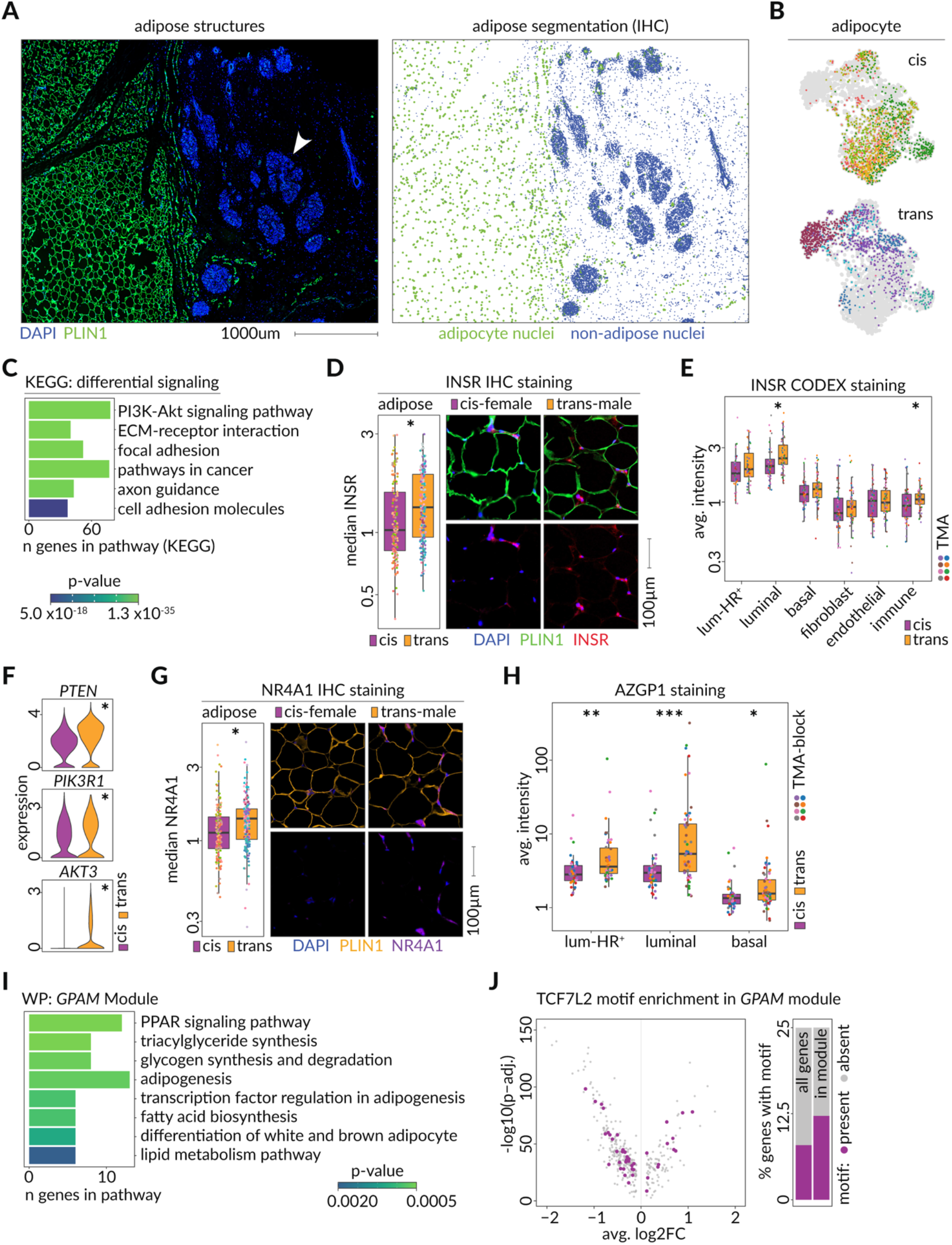
Regulatory and spatial transition of adipocytes upon androgen hormone replacement therapy. **A)** Left microscopic image stained against nuclei (DAPI; blue) and adipocytes (PLIN1; green) and right shows the resulting nuclei segmentation. **B)** UMAP of adipocytes showing cis-female (top) and trans-male (bottom) cells colored by sample origin. **C)** Pathway enrichment (KEGG) of matched ligands and receptors that are differentially expressed between trans-male and cis-female samples in all mammary cell types. **D)** Boxplots show median INSR staining intensity in IHC scan-regions of cis-female (purple) and trans-male (orange) samples (p-value, Wilcoxon: 0.00043). Right shows IHC micrographs stained against nuclei (DAPI; blue), adipocytes (PLIN1; green), and INSR (red) in cis-female (left) and trans-male (right) tissues. **E)** Boxplots show the average staining intensity of INSR in cis-female (purple) and trans-male (orange) cell types in tissue regions of the CODEX microarray. Dot colors indicate the origin TMA of the shown regions (p-values, Wilcoxon: luminal = 0.0093, immune = 0.014). **F)** Boxplots show median NR4A1 staining intensity in IHC scan-regions of cis-female (purple) and trans-male (orange) samples (p-value, Wilcoxon: 7.5 x 10^-5^). Right shows IHC micrographs stained against nuclei (DAPI; blue), adipocytes (PLIN1; orange), and NR4A1 (purple) in cis-female (left) and trans-male (right) tissues. **G)** Violin plots show the expression of *PTEN* (top), *PIK3R1* (middle), and *AKT3* (bottom) in cis-female (purple) and trans-male (orange) adipocytes (p-values, MAST: *PTEN* = 9.04 x 10^-96^, *PIK3R1* = 5.75 x 10^-38^, *AKT3* = 1.41 x 10^-18^). **H)** Boxplots show the average staining intensity of AZGP1 in cis-female (purple) and trans-male (orange) epithelial cell types in tissue regions of the CODEX microarray. Dot colors indicate the origin TMA of the shown regions (p-values, Wilcoxon: luminal-HR+ = 0.0011, luminal = 0.00047, basal = 0.011). **I)** Pathway enrichment (WP) of *GPAM* co-expression module genes that are differentially expressed between trans-male and cis-female adipocytes. **J)** Volcano plot shows the average log_2_ fold change and –log_10_ adjusted p-value for differential expression of genes within the *GPAM* co-expression module among the trans-male and cis-female adipocytes. Purple data points indicate genes with a chromatin accessibility peak overlapping the TCF7L2 transcription factor sequence motif (CisBP TCF7L2_762) match. Barplots show the fraction of all genes (left) or genes within *GPAM* module (right) which contain a chromatin accessibility peak overlapping the TCF7L2 transcription factor sequence motif (purple).

**Supplementary Figure S13:**
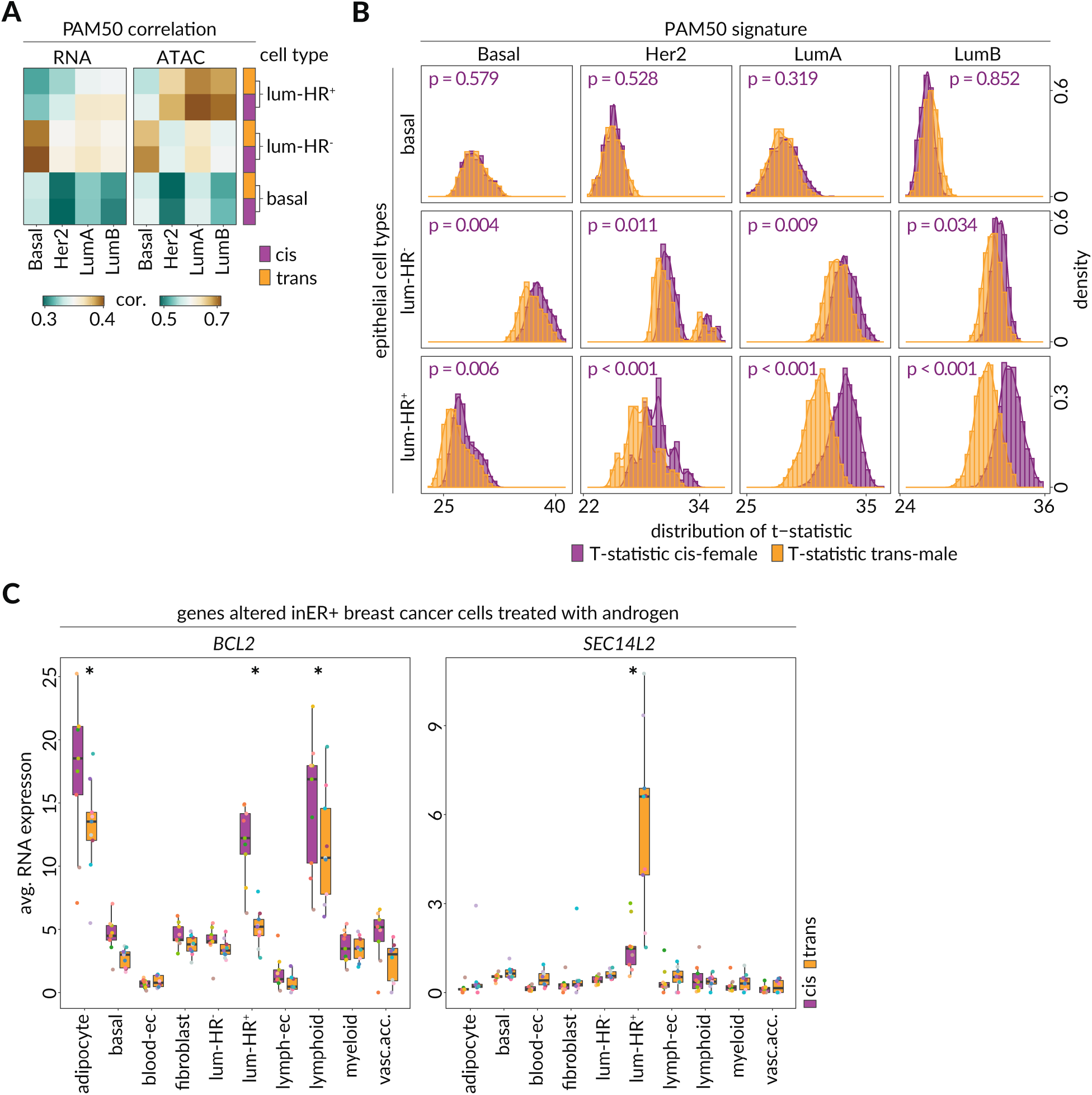
Association of breast epithelial cell types with PAM50 signatures. **A)** Heatmaps show Pearson correlation of log_2_(FPKM + 1) of gene expression (left) and chromatin accessibility FPM (right) among one of the epithelial cell types of cis-female (purple) and trans-male (orange) samples compared to one of the four major PAM50 subtypes of TCGA breast cancer datasets. **B)** Histograms show the kernel density estimate for the distribution of t-statistic in 1,000 permutations comparing the t-statistic of a linear model for association of a randomly sampled subset of TCGA breast cancer samples with a randomly sampled subset of a cis-female (purple) epithelial cells and a randomly sampled subset of trans-male (orange) epithelial cells (rows). P-values indicate the fraction of 1,000 permutations where a subset of trans-male epithelial cells had a higher t-statistic than a subset of cis-female epithelial cells. **C)** Average RNA-expression of AR-targeted oncogene *BCL2* and tumor suppressor *SEC14L2* (discussed in Hickey et al, Nat Med, 2021) in trans-male (orange) and cis-female (purple) samples. (p-values, MAST, *BCL2*: adipocyte = 1.72 x 10^-34^, lum-HR^+^ < 1.72 x 10^-34^, lymphoid = 7.28 x 10^-22^, *SEC14L2*: lum-HR^+^ < 2.22 x 10^-16^).

## Materials and Methods

### Sample collection

Fresh human breast tissue specimens from elective mammary surgeries in cisgender women and subcutaneous mastectomies in transgender men were collected at time of surgery. Tissues were sampled from the central breast, inferior and deep to the nipple (6 o’clock), placed in Dulbecco’s Modified Eagle’s Medium (DMEM, Corning) and processed shortly after. Tissues were washed 3x with DMEM, and any large pieces of fat were grossly removed. Tissue was then cut into 2-3 mm^3^ pieces before being directly stored at −80 °C. Where available, the Cedars-Sinai Biobank collected pieces of tissue from the left and right breast and fixed them with 10% formalin for subsequent paraffin embedding (FFPE). This study was approved by the Institutional Review Board (IRB) of Cedars-Sinai Medical Center.

### Preparation of RNA expression libraries

All RNA-expression libraries were prepared with Chromium Single Cell 3’ (v3) Reagent Kits by 10X Genomics. To assure minimal handling time while reducing batch effects, samples were processed in batches of 3-4 (with each batch containing trans-male and cis-female samples). All steps until library generation were carried out on ice and in pre-cooled instruments. Lysis was done with Nuclei EZ lysis buffer (Sigma) and subsequent steps were carried out in a custom wash buffer (10 mM Tris, 146 mM NaCl, 1 mM CaCl2, 21 mM MgCl2) that was freshly supplemented with 40 U/mL of RNAse inhibitor (Thermo Fisher) and 2% of Molecular Biology Grade BSA (New England Biolabs). About 250 mg of cryopreserved tissue was placed on wet ice and cut with a scalpel into rice grain sized pieces while still frozen. The tissue was then transferred into a chilled 7 mL dounce tissue grinder (Sigma) containing 3 mL of 20% Nuclei EZ lysis buffer (diluted 1:5 with wash buffer). Tissue was soaked on ice for 3 min with occasional pipetting with a wide bore tip. Lysate was homogenized, first with pestle marked “A” (coarse) and then “B” (fine), for 10 complete strokes and for no longer than 5 min. Nuclei suspension was then filtered through a 70 μm filter (Miltenyi) and the lysis buffer was quenched with 9 mL of wash buffer. Filtrate was then spun down in a swing bucket rotor for 8 min at 850 g, and pellet was resuspended in 1.2 mL of wash buffer. Suspension was filtered through a 20 μm filter (Miltenyi), spun down again and final pellet was resuspended in 400 μL of wash buffer supplemented with 2.5 μg/mL DAPI. Solution was immediately loaded onto a FACSAria III Cell Sorter (BD Biosciences) equipped with a 70 μm nozzle (liquid output for this setup was previously measured to be 1.9 nL per event), and a gate around single nuclei was determined using DAPI and side-scatter signals. 22,000 events (∼41.8 μL) were sorted directly into a 96-well round bottom plate harboring RT-buffer (10X Genomics), which was prepared without adding RT-Enzyme (total of 25.1 μL). After sorting was completed, RT-Enzyme (8.3 μL) was added, and nuclei suspension was immediately loaded onto a 10X Chromium controller. Following library preparation was performed according to the Chromium Single Cell 3’ Reagent Kits User Guide (v3 Chemistry) with an assumed input of 10,000 cells. The cDNA amplification was carried out with two extra PCR cycles (13 total); all other steps were kept unaltered. Quality, amount and size distribution of the final libraries was assessed on a BioAnalyzer (Agilent).

### Preparation of single nuclei ATAC libraries

All snATAC libraries were prepared with Chromium Single Cell ATAC Reagent Kits (v1.1) by 10X Genomics. Nuclei extraction method was identical to RNA-expression workflow (see above) with the following buffers (prepared according to 10X Genomics demonstrated protocol: CG000212, Rev.B). Lysis Buffer (10 mM Tris-HCl pH 7.4, 10 mM NaCl, 3 mM MgCl2, 0.01% Tween-20, 0.01% Nonidet P40 Substitute, 0.001% Digitonin, 1% BSA), Wash Buffer (10 mM Tris-HCl pH 7.4, 10 mM NaCl, 3 mM MgCl2, 0.1% Tween-20, 1% BSA). After washing and 20 μm filtration (see RNA-expression workflow above), nuclei were spun down and the final pellet was resuspended in 400 μL of wash buffer, supplemented with 0.5 μg/mL 7-AAD (BioLegend). Solution was immediately loaded onto a FACSAria III Cell Sorter (BD Biosciences) equipped with a 70 μm nozzle and a gate around single nuclei was determined using 7-AAD and side-scatter signals. All available nuclei (on average ∼350.000 events) were sorted directly into a 1.5 mL protein lo-bind tube (Eppendorf) containing 100 μL diluted nuclei buffer (10X Genomics Single Cell ATAC Reagent Kit). The sort volume was calculated, and the sorted nuclei were supplemented with 20x diluted nuclei buffer to a final concentration of 1x. Nuclei were spun down at 850 g for 8 min, supernatant was carefully removed as much as possible and the final pellet was resuspended in 10 μL diluted nuclei buffer. 5 μL of nuclei suspension was mixed with 5 μL of 5 μg/mL DAPI and counted on a hemocytometer to determine loading concentration. If necessary, the remaining nuclei solution was diluted to 3000 nuclei/μL and used immediately in the transposition reaction. Following library preparation was performed according to the Chromium Single Cell ATAC Reagent Kits User Guide (v1.1 Chemistry). Quality, amount and size distribution of the final libraries was assessed on a BioAnalyzer (Agilent).

### Sequencing

Single nuclei RNA expression libraries were sequenced according to 10X Genomics recommended read lengths (read1 = 28bp, read2 = 91bp) on a NovaSeq6000 (Illumina) to an average of ∼40.000 reads per nucleus, resulting in an average sequencing saturation of ∼75% (as reported by 10X Genomics CellRanger v6.0.1). Single nuclei ATAC libraries were sequenced according to 10X Genomics recommended read lengths (read1 = 50bp, read2 = 50bp) on a NovaSeq6000 (Illumina), to an average of ∼32.000 reads per nucleus, resulting in an average of ∼7000 fragments per cell (as reported by 10X Genomics CellRanger ATAC v1.2.0).

### Preprocessing of single nuclei RNA expression data

Fastq files were processed using 10X Genomics CellRanger v6.0.1 and aligned to the human reference genome *“refdata-gex-GRCh38-2020-A”* provided by 10X Genomics. In order to account for increased amounts of pre-mRNA captured in single nuclei RNA sequencing, CellRanger was run using the option *—include-introns*. Resulting count matrices were further processed in Seurat^116^. Thresholds for maximum fraction of mitochondrial genes and number of UMIs for each nucleus were set to 2.5% and 20.000 respectively. Barcodes that likely contained doublets were detected and removed with scDblFinder (v1.6.0)^116^. Further doublet-enriched and low UMI (median UMI count <1000) clusters that emerged while sub-clustering each individual cell type were also removed.

### Dimensional reduction, cell type and subcluster identification in snRNA-seq data

All samples were integrated into a single dataset using the standard Seurat workflow (variable features = 5000, principle components = 50, louvain resolution = 0.05), batch-corrected with Harmony v0.1.0, and projected into two dimensional space using uniform manifold approximation and projection (UMAP)^119, 120^. Identification of the main cell types was done by using canonical marker genes (Fig. S1D) and further confirmed through existing gene modules in case of the epithelial cells (Lim et. al., 2009)^121^.

Each individual cell type was then extracted into a separate dataset for further classification of subclusters. Lymphoid and myeloid subtypes were determined by using canonical immune markers (Fig. S11A). Blood endothelial subclusters were determined via conserved marker modules from *Table S5*, (Kalucka et. al., 2020) (Fig. S10B) and pericytes were distinguished from vascular smooth muscle cells through expression of ACTA2 and PDGFRB (Fig. S10C)^56^. Fibroblast subclusters were labeled to reflect the function of their top marker genes.

### Differential gene expression, pathway enrichment, co-expression module generation, gene module scoring and receptor ligand interactions in snRNA-seq data

Differential gene expression analysis on snRNA-seq data was done on log-transformed counts using MAST, and filtered for FDR < 0.05^122^. Curated gene sets were obtained from MSigDB v7.2 from the *“h:hallmark”* and *“c2:curated”* gene set collections and filtered for Reactome, PID, WikiPathways, Biocarta, KEGG and Hallmark as providers, as well as a minimum set size of 10 genes^123^. Nuclei were scored for gene modules according to scanpy’s *score_genes* tool, using default settings v1.4.5.1 and full results of these analyses can be found in Table 5 ^124^. Pathway enrichment of gene modules was carried out via the R client for g:Profiler v0.2.1 and full results for the relevant plots can be found in Table 6^125^. Gene regulatory networks were generated using GRNboost2 with a list of 1839 transcription factors and pathway anchors *(hs_hgnc_tfs)* provided in the resources of pySCENIC^126, 127^. From the resulting TF-target association table, highly correlated target genes (>95^th^ percentile of importance) of a transcription factor were selected to form a co-expression module. A curated list of ligand-receptor interaction pairs was taken from Cabello-Aguilar et. al., 2020, and filtered for pairs with a PMID reference^128^.

### GTEx breast tissue and sex bias analysis

Gene RNA-expression data in other tissues was acquired through the Genotype-Tissue Expression Project (version 8) (GTEx) which was supported by the <u>Common Fund</u> of the Office of the Director of the National Institutes of Health, and by NCI, NHGRI, NHLBI, NIDA, NIMH, and NINDS^129^.

Clustering and dimensionality reduction of GTEx breast samples was done with selected tools of the Seurat preprocessing workflow (variable features = 2000, principle components = 50, louvain resolution = 0.2) (Fig. S6). The resulting dataset was scored for the top 100 marker genes of the epithelial and vasculature groups, as well as the top 100 marker genes of the adipocyte and fibroblast cell types taken from our snRNA-seq data (Fig. S6C, D). The resulting module scores were used to classify the breast samples into epithelial, vasculature and adipose enriched subsets. Since the fibroblast score did not highlight a distinctive cluster, we chose the 50^th^ percentile of the fibroblast score for this subset. We used the GTEx table of gene counts (GTEx Analysis 2017-06-05 v8 RNASeQCv1.1.9) to identify the differentially expressed genes among cis-male and cis-female breast samples using DESeq2 (v1.32.0)^130^. We then compared these genes to the corresponding significant differentially expressed genes in snRNA-seq data.

For the CUX2 and sex bias analysis, median transcript TPM values were calculated using the aggregated tissue classifier *“SMTS”* of the GTEx “Sample Attributes” metadata and sex bias effect sizes were used as provided by GTEx *“GTEx_Analysis_ v8_sbgenes.tar.gz”.* In summary: sex-biased gene expression statistics for GTEx v8 tissues present in both sexes are derived from across-tissue meta-analysis with MASH (Urbut et al. 2019), based on per-tissue sex effect size and corresponding standard error values calculated with voom-limma (Law et al. 2014)^131, 132^.

### Inferring trajectory and diffusion maps from snRNA-seq data

To prepare input for RNA-velocity analysis, the BAM files generated by the CellRanger snRNA-seq workflow were processed into loom files containing spliced and unspliced abundances using velocyto v0.17.17^133^. Here, the same 10X Genomics genome annotation file used in CellRanger *“refdata-gex-GRCh38-2020-A”* was utilized alongside the GRCh38 repeat mask file from UCSC genome browser ^134^. The loom files were combined and cells that passed QC in previous gene expression analysis were extracted from the output. RNA-velocity analysis was done on each cell type separately using the standard workflow of the scVelo package v0.2.3 and cellrank v1.3^135, 136^. Moments were calculated using the batch-corrected first 30 harmony principal components, and velocity was estimated using the top 2,000 most variant genes and the dynamical model. For CellRank analysis, we determined the terminal states using the prior information of having two terminal states within each cell type (corresponding to cis-female and trans-male populations), weight connectivity of 0.2, and the Monte Carlo average of randomly sampled velocity vectors. We used the default parameters to identify the initial state.

To extract the diffusion maps, we used the Bioconductor package destiny v3.0.1^137^. We used snRNA-seq data of each cell type with the default parameters of the *DiffusionMap* function.

### Preprocessing and integration of single nuclei ATAC data

We used 10X Genomics CellRanger ATAC (v1.2.0) to align the fastq files to the reference genome *“refdata-cellranger-atac-GRCh38-1.2.0”* and obtained the fragment and barcode annotation file for each sample. The resulting 500 bp resolution data was further processed using ArchR v1.0.0 to remove doublets and correct for batch effects using Harmony^138^. Reduced representation of the batch-effect–corrected tile matrix was calculated with iterative latent semantic indexing, and visualized with UMAP. We then used ArchR to integrate snATAC-seq data with snRNA-seq data by correlating gene activity scores of each snATAC-seq cluster with the transcriptome of each snRNA-seq cluster (Fig. S1D).

### Footprint analysis, enhancer mapping and differentially accessible peak identification

All transcription factor binding motifs were provided by the Catalog of Inferred Sequence Binding Preferences (CisBP)^139^. We used the motif footprint feature of ArchR to investigate the enrichment of each motif among the peaks of each cell type or gender identity while adjusting for the Tn5 bias by dividing the signal to Tn5 bias. ArchR investigates the correlation of each peak with the expression levels of a gene, where we used a Pearson correlation cutoff of 0.2 to identify peak-gene relationships. We used ArchR’s implemented Wilcoxon test to identify differentially accessible peaks adjusting for transcription start site enrichment (TSS) and number of fragments in log10.

### Motif enrichment in cell types and single nuclei

To calculate motif enrichment for each cell type on basis of gender identity, we compared the enrichment of each motif in the foreground (trans-male nuclei) versus the background (cis-female nuclei). We performed a motif enrichment analysis through ArchR and using CisBP motif database. The motif enrichment analysis investigates the peaks accessible within the group that contains the sequence motif of interest. It compares the fraction of peaks containing the motif in the foreground versus the background and calculates a one-sided Fisher’s exact test. For the top motifs passing the cutoff adjusted p-value < 0.05, we visualized the odds ratio of the enrichment for each cell type. Motif enrichment on single nuclei basis was done using chromVAR, which calculates z-scores that indicate the enrichment of each sequence motif in each nucleus relative to other nuclei^140^.

### HiC data analysis

We downloaded two publicly available HiC datasets for MCF-10A (Accession: GSE98552) and PANC-1 (ENCODE Accession: ENCLB951HSJ)^141, 142^. MCF-10A data had already been processed using HiC-Pro v2.8.1, GRCh38 genome assembly, and 40,000 bp genomic bins^143^. We used the same software and pipeline to process PANC-1 datasets. We divided the ICE-normalized genomic bin contact values to the maximum value of the chromosome to adjust for the potential variations introduced by library size and experimental protocol. We visualized contact of genomic bins overlapping the RYR2 gene or its enhancers that passed the 75^th^ percentile after pooling data of MCF-10A and PANC-1 around RYR2.

### High-resolution peak calling and motif filtering

In addition to the peak set generated through ArchR, we also generated pseudo-bulk BAM files by extracting the reads corresponding to each cell type and sample type (total of 20 BAM files). For each pseudo-bulk BAM file, we used MACS2 peak calling with parameters *-f BAMPE -g 2.7e9 --nomodel --SPMR --bdg*^144^. We then used the generated *NarrowPeak* files for identifying motifs that are within an absolute distance of 200 bp from the summit of the peaks.

### Predicting dysregulation direction of AR-regulated genes with Random Forest

We identified 603 genes upregulated in trans-male luminal-HR^+^ cells (adjusted p-value < 0.01 and log_2_ FC > 0.25) and 652 genes downregulated (adjusted p-value < 0.01 and logFC < −0.25). To consider a gene an AR-regulated gene, we required an ARE containing peak whose accessibility correlates with the gene’s expression (Pearson r > 0.2) as identified by ArchR (see above). AR upregulated 309/603 genes and downregulated 244/652 genes. ArchR identified 1,080 500 bp peaks which correlate with the expression of these 553 genes. Using pseudo-bulk BAM files, we identified 1,427 narrow peaks corresponding to these 500 bp genomic regions. These narrow peaks ranged from 141 bp to 3,011 bp with a median of 792 bp. We annotated each peak according to the chromatin accessibility signal at the exact motif position in the peak in luminal-HR^+^ trans-male cells for motifs of 189 expressed transcription factors. We labeled these peaks according to their mapping to downregulated genes or upregulated genes. We randomly selected 80% of these peaks as the training set and the remaining 20% as our validation set. We trained a random forest with 500 trees with each tree sampling 100 of the 189 motifs to predict if the gene was downregulated or upregulated. We used the R package ‘randomForest’ (v4.6-14). We allowed each classification tree to reach maximum purity. The model was able to predict downregulated genes with an area under the receiver-operating characteristic curve (auROC) of 0.825 and area under the precision-recall curve (auPR) of 0.741. We used mean decrease in Gini index as the measure of variable importance.

### Enrichment of transcription factors in pathways

To investigate if a transcription factor regulates a particular pathway, we identified the number of genes of that pathway which have a chromatin accessibility peak containing the motif of that transcription factor. We also required the chromatin accessibility peak to correlate positively with the expression of the gene (Pearson r > 0.2). We compared the fraction of pathway’s genes containing the motif in their peaks to a background of all the genes using a one-sided Fisher’s exact test.

### Identifying active transcription factors

Sequence motifs corresponding to the same family of transcription factors often show similar binding sites. To identify the active transcription factors, we used ArchR’s correlation of a motif’s z-score versus the expression of its corresponding transcription factor. For the specific case of nuclear receptors in luminal-HR^+^ cells, we built all of the possible simple linear models which predict a nuclear receptor’s motif’s z-score given the expression of all other nuclear receptors (one at a time) or a background transcription factor (i.e. *PPARG*). To be able to compare the effect sizes, we standardized both the motif’s z-score as well as the transcription factor’s expression. We compared the linear model effect size, p-value, and R^2^.

### Immunohistochemistry

Immunohistochemistry was performed on sections taken from FFPE blocks that were collected by the Cedars Sinai Biobank at time of surgery. Briefly, fixed sections were incubated in 60 °C for 25 minutes to remove excess paraffin and then immediately deparaffinized and rehydrated. Antigen retrieval was performed using an “Instant Pot® Duo™” pressure cooker and 1x Universal HIER buffer (Abcam, cat: ab208572). Background fluorescence was quenched by photobleaching for 1.5h in bleaching solution according to (Du et al.)^145^. Sections were then blocked in protein blocking buffer (Abcam, cat: ab64226) for 1 h at room temperature, washed and then incubated with primary antibodies at 4 °C overnight. The primary antibodies used were as follows (all dilutions were performed with protein blocking buffer): KRT8 (Sigma-Aldrich, cat: MABT329, clone TROMA-1, 1:100), PLIN1 (Thermo-Fisher Scientific, cat: PA5-81240, rabbit polyclonal, 1:500), TCF4 (Sigma-Aldrich, cat: 05-511, clone 6H5-3, 1:100), Nur77-AF488 preconjugated (Thermo-Fisher Scientific, cat: 53-5965-82, clone 12.14, 1:500), INSR (Abcam, cat: ab90940, clone MM0414-2A12, 1:100), CD45 (Abcam, cat: 40763, clone EP322Y, 1:200), CD68 (Thermo-Fisher Scientific, cat: MA5-13324, clone KP1, 1:100), and CD3e-AF488 custom conjugated (Abcam, cat: 271850, clone EP449E, 1:200).

Sections were then washed and incubated with the appropriate fluorophore-conjugated secondary antibodies at room temperature for 1 hour. Secondary antibodies used were as follows (all dilutions were performed with protein blocking buffer): Goat anti-rabbit IgG AF488 (Thermo-Fisher Scientific, cat: A11008, 1:500), Goat anti-rabbit IgG AF568 (Thermo-Fisher Scientific, cat: A11011, 1:500), Donkey anti-mouse IgG AF568 (Thermo-Fisher Scientific, cat: A10037, 1:500), and Goat anti-rat IgG Cy5 (Thermo-Fisher Scientific, cat: A10525, 1:500).

Sections were finally Washed 3 times with 1X PBST (1X Phosphate Buffered Saline with 0.1% Tween-20) for 3-5 minutes at room temperature, mounted with Vectashield containing DAPI (Vector Laboratories, cat: H-1200), and imaged using a Zeiss Axio Scan.Z1.

Automated imaging was set up in Zeiss ZEN pro v3.1. Due to large tissue sizes, the slides were scanned without z-stack and divided into ∼12-25 regions with individual focus maps. CZI format images were read with the AICSImageIO package (version 3.3.7)^146^. Nuclei from the DAPI channel of each image were segmented with StarDist, then the average staining intensity per nucleus was tabulated^147, 148^.

### Co-detection by indexing (CODEX) of tissue microarrays

Tissue microarrays (TMA) were prepared from 2 mm punches taken from left and right breast representative FFPE blocks where available. To ensure the capture of diverse tissue sections that include epithelial structures, H&E stains were used to pre-annotate regions of interest, which were then transferred onto TMA paraffin blocks. Each (4×4) block contained four regions from four patients (see Fig. S1C) and each patient was represented with eight regions across two separate TMA-blocks, resulting in eight TMAs total.

Sections from each of these 8 TMAs were then collected onto poly-L-lysine-coated coverslips, which were prepared according to the Akoya Biosciences CODEX protocol. Similar to the IHC protocol (see above), sections were incubated in 60 °C for 25 minutes, then deparaffinized and rehydrated. Following antigen retrieval, sections were then quenched for autofluorescence using a protocol adapted from (Du et al.)^145^. Subsequently, sections were stained and imaged according to the Akoya Biosciences CODEX protocol. Details regarding primary antibodies and imaging conditions can be found in Supplementary Table 7. Imaging was performed using a Leica DMi8 equipped with a 20x objective, Lumencor SOLA SE U-nIR LED, and Hamamatsu Orca Flash 4.0 v3.

Primary antibodies were initially screened by performing standard IHC (as above) on FFPE tissue sections to verify positive staining. Primary antibodies were then conjugated to their corresponding barcodes according to the Akoya Biosciences CODEX antibody conjugation protocol. Conjugated antibodies were then titrated by performing CODEX staining on a TMA section using the full panel diluted at either 50x, 100x, or 200x. The dilution that resulted in the optimal signal-to-noise ratio was determined for each antibody individually. The final dilutions obtained from this titration can be found in Supplementary Table 7.

### CODEX data pre-processing

Raw images of TMA-regions from CODEX experiments were pre-processed with a custom workflow where five preprocessing operations were applied in this order: extended depth of field (EDOF), shading correction, cycle alignment, background subtraction and tile stitching, described briefly here.

1: An EDOF image was produced from the z-stack for each tile where each position is taken from the z-plane most in focus. 2: The BaSiC method of optical shading correction was applied to each channel of each imaging cycle^149^. 3: An image registration transformation was estimated between the first cycle DAPI channel and the DAPI of each subsequent cycle. For each cycle, the registration parameters were saved and applied to all other channels from the same cycle. 4: Blank cycles were used to subtract background from each channel. 5: Finally, neighboring tiles were stitched by applying a registration between the overlapping areas between two tiles. First, the two tiles with the best naive overlap were stitched by applying the appropriate registration shift to one of the tiles. Stitching then proceeded with the next two most nearly aligned tiles, until all tiles were merged. Since each cycle was previously aligned to the first cycle’s DAPI channel, the registrations used for tile stitching were estimated once on the first DAPI and reused for subsequent channels and cycles.

The assembled DAPI images for each TMA region were then visually inspected and any remaining grossly out-of-focus areas due to low tissue adherence were removed from further analysis.

To obtain nuclear segmentations, we applied a pre-trained StarDist model to the first cycle DAPI image^147, 148^. The model weights of the 2D 2018 Data Science Bowl model released by the original StarDist authors were fine-tuned using a training set of nuclei imaged on our CODEX platform. A whole-cell or “cytoplasmic” segmentation was obtained by expanding the nuclear segmentation area by morphological dilation, without introducing overlaps in adjacent nuclei. The average intensities under each nuclear mask and cytoplasmic mask were extracted for each cell to be used for cell type assignment.

### CODEX data analysis

Nuclei and cytoplasmic staining intensities of all data points passing segmentation were merged into a single unfiltered dataset, followed by batch correction, dimensional reduction and clustering with harmony and Seurat. The dataset was then cleaned from over-segmentation (i.e. blood clots, very dim regions, damaged tissue) by visually inspecting sub clusters and removing data points that were not represented by a clearly visible nuclear DAPI signal. Cell classes were defined according to their marker staining and the resulting filtered dataset was then scaled according to the 1st and 3rd quartiles using RobustScaler across each region, producing the intensity values shown in log scale^150^. When calculating nuclear vs. cytoplasmic staining, raw signal values were used. Where region averages of these values are shown, only regions with a minimum of 10 cells of the discussed cell type are shown, unless otherwise stated.

### Epithelial neighborhood and cell-cell distance analysis

For all analyses including cell-cell adjacencies (CODEX and IHC), we calculated pairwise Euclidean distances based on the nuclei centroids using the package rdist v0.0.5. To assess the composition of cell types that are closely associated with the epithelium we first detected spatial clusters of epithelial cells (cells in cluster > 35) using affinity propagation from the APCluster v1.4.9 package^151^. To define the epithelial neighborhood populations, we retained all cells within 100px (∼32.5µm) distance from these epithelial cells.

### Segmentation of acini and assessment of epithelial smooth muscle layer

To study the staining patterns around epithelial cells in detail, we applied morphological operations to specifically isolate individual acini structures within the CODEX tissue microarray images (Fig. S8C). First, masks of the KRT8 and KRT23 stains per TMA region were obtained by multi otsu thresholding (scikit-image), and imposing a minimum intensity of 100 in order to avoid false-positive epithelial segmentations^152^. An ACTA2 mask was obtained similarly by multi otsu thresholding, but without the minimum intensity requirement. Morphological dilation was applied to each mask. Then, the KRT8 and KRT23 masks were added together to form a pan-luminal mask, and the ACTA2 mask was subtracted from this pan-luminal mask to divide acini into individual instances. Individual putative acini were labeled by connected components, dilated to fill extraneous holes, filtered for a minimum area of 200, then individually analyzed. Segmented nuclei underneath each acini mask were gathered to tally the cell type composition, and the 15-pixel wide border surrounding the acini was considered the border area. The percentage of the acini border that was positive for ACTA2 above the predetermined threshold was tallied.

### Image analysis of adipocyte vacuole area

To study the morphology of adipocytes we used IHC staining of PLIN1 to segment individual adipocyte vacuoles and profile their size. To avoid abundant small-scale staining artifacts, PLIN1 IHC images were analyzed at 1/5th the original resolution. First, the PLIN1 signal was capped at the 90^th^ percentile intensity value and thresholded using Ostu’s method (scikit-image v0.19.1). To join small gaps in the PLIN1 mask we applied morphological dilation (disk structuring element radius 2) and closing (disk structuring element radius 5). A label image was created from the inverse of the PLIN1 mask, where segmented regions represented regions of PLIN1^-^ area that were completely enclosed by PLIN1^+^ staining. We next filtered these putative vacuoles to exclude artifacts including small regions, and regions corresponding to multiple vacuoles erroneously connected because of e.g. broken membrane. To filter putative vacuoles, we calculated the area, and convex area (area of a convex hull fit around the actual label) and took their ratio. We reasoned that true positive vacuole segmentations would have an area-to-convex-area ratio (Rac) near 1 and applied a threshold of Rac > 0.85. Similarly, we reasoned that erroneously connected vacuoles, or over-segmentations, would be outliers on the distribution of area, and applied a lower threshold determined by Otsu’s method of all vacuole areas exceeding a predetermined absolute minimum of 101.25 pixels, and a fixed upper threshold of ∼103.7 pixels. Following these filters, we noted that the PLIN1-INSR experiment had consistently more coverage of adipose vacuoles than others, and restricted further analysis to these images. Each image scene containing more than 1,000 adipocyte nuclei (determined by DAPI segmentation and PLIN1 threshold) was summarized by the median area of observed vacuoles.

## Contributions and Acknowledgements

F.R., E.C.R, X.C. and S.R.V.K. conceived of the study. E.C.R. performed the gender-affirming surgeries and A.E.G. performed the cisgender women surgeries, with both contributing to the tissue collection process along with Y.Q and T.Y.L. and X.C. F.R., Q.Y, A.M., T.Y.L. performed the nuclei isolation and F.R. prepared all sequencing libraries. F.R., M.K., N.I., B.M. performed all computational analysis with B.W., H.G. and S.R.V.K. supervising. F.R., M.K., N.I., B.M., S.D., B.W., H.G., E.C.R., X.C. and S.R.V.K. interpreted the data and wrote the manuscript. We would like to thank Dr. Benjamin Hopkins and Dr. Camilla Dos Santos for helpful discussions about this dataset as well as the Cedars-Sinai Medical Center Biobank and Translational Genomics Core for their helpful support of this project.

## Data Availability

All raw sequencing data reported in this manuscript will be uploaded to a suitable public repository prior to publication.

## Code Availability

All computing code described in this manuscript will be uploaded to a suitable public repository prior to publication.

## Notes

### Competing Interest Statement

The authors have declared no competing interest.

